# Heterogeneous disease-propagating stem cells in juvenile myelomonocytic leukemia

**DOI:** 10.1101/628479

**Authors:** Eleni Louka, Benjamin Povinelli, Alba Rodriguez Meira, Gemma Buck, Neil Ashley, Angela Hamblin, Christopher A.G. Booth, Nikolaos Sousos, Anindita Roy, Natalina Elliott, Deena Iskander, Josu de la Fuente, Nicholas Fordham, Sorcha O’Byrne, Sarah Inglott, Anupama Rao, Irene Roberts, Adam J. Mead

## Abstract

Juvenile Myelomonocytic Leukemia (JMML) is a poor prognosis childhood leukemia usually caused by germline or somatic RAS-activating mutations. The cellular hierarchy in JMML is poorly characterized, including the identity of leukemia stem cells (LSCs). FACS and single-cell RNA-sequencing reveal marked heterogeneity of JMML hematopoietic stem/progenitor cells (HSPCs), including an aberrant Lin-CD34+CD38-CD90+CD45RA+ population. Single-cell HSPC index-sorting and clonogenic assays show that (1) all somatic mutations can be backtracked to the phenotypic HSC compartment with *RAS*-activating mutations as a “first hit”, (2) mutations are acquired with both linear and branching patterns of clonal evolution and (3) mutant HSPCs are present after allogeneic HSC transplant before molecular/clinical evidence of relapse. Stem cell assays reveal inter-patient heterogeneity of JMML-LSCs which are present in, but not confined to, the phenotypic HSC compartment. RNA-sequencing of JMML-LSCs reveals upregulation of stem cell and fetal genes (*HLF, MEIS1, CNN3*, *VNN2*, *HMGA2*) and candidate therapeutic targets/biomarkers (*MTOR, SLC2A1*, *CD96*) paving the way for LSC-directed disease monitoring and therapy in this disease.

## Introduction

Juvenile myelomonocytic leukemia (JMML) is an aggressive subtype of childhood myelodysplastic syndrome (MDS), usually presenting in the first 5 years of life, that is characterized by abnormal proliferation of dysplastic cells of the monocytic and granulocytic lineages (*1*). Although JMML is a rare childhood cancer, it has several key features which make it an important paradigm. In particular, it is definitively caused by RAS-activating mutations, typically in *PTPN11*, *KRAS*, *NRAS*, *CBL* or *NF1* in >90% of cases and the molecular landscape appears otherwise relatively simple with a low number of somatic mutations in comparison with other malignancies (*2, 3*). Some patients show evidence of clonal evolution, with acquisition of monosomy 7 (*1*) or secondary somatic mutations of *SETBP1, ASXL1*, *EZH2* and other genes which are associated with a worse prognosis (*2, 3*). The only curative therapy for JMML is allogeneic hematopoietic stem cell transplantation (HSCT), however, relapse rates are high (*1*) indicating a failure to eradicate the disease-propagating cells in this condition.

The presence of distinct populations of rare disease-propagating cancer stem cells (CSC) has been demonstrated in some cancers, and this is a crucial step towards understanding cellular pathways of disease relapse (*4*). In adults with chronic myeloid neoplasms, including myeloproliferative neoplasms (MPN) (*5*), chronic myeloid leukemia (*6*) and MDS (*7*), rare and distinct CSCs have been identified which share phenotypic features with normal HSCs. However, in acute myeloid leukemia (AML), leukemia stem cells (LSCs) are more heterogeneous and, although the disease may originate in HSCs, various different progenitor cell populations are transformed and are responsible for disease propagation (*8*). Furthermore, in childhood acute lymphoblastic leukemia (ALL), blast cells across all stages of differentiation have LSC properties (*9*). In view of this marked heterogeneity of CSCs/LSCs between different liquid tumors, it is crucial to properly characterize the CSCs in specific disease entities, particularly where the prognosis is poor and targeted therapy remains elusive. Indeed, for JMML, while the genetic basis is now well described, very little is known about the cellular hierarchy, including the identity of cells that propagate the disease and cause relapse.

## Results

### Single cell phenotypic, functional and molecular analysis reveals heterogeneity of hematopoietic stem/progenitor cells (HSPCs) in JMML

In order to characterize LSCs in JMML, we established a national prospective study and collected serial bone marrow (BM) and peripheral blood (PB) samples from a cohort of 16 patients with JMML for phenotypic, functional and molecular analysis of HSPC pre- and post-HSCT (Table S1; Figure 1A). At diagnosis, the phenotype of early (CD38 negative) HSPC was markedly disrupted in JMML (Figure 1B and 1C). Although numbers of phenotypic HSCs (Lin-CD34+CD38-CD90+CD45RA-) were normal, multipotent progenitors (MPP; Lin-CD34+CD38-CD90-CD45RA-), including lymphoid primed MPPs (LMPP; Lin-CD34+CD38-CD90-CD45RA+) were reduced in JMML in comparison with normal pediatric BM (Figure 1B and 1C). A number of patients (8/14 analyzed) also showed the presence of an aberrant Lin-CD34+CD38-CD90+CD45RA+ (+/+) population of cells. In contrast, the CD38+ myeloid progenitor compartment showed an apparently normal frequency of phenotypically defined common myeloid progenitors (CMP; Lin-CD34+CD38+CD123+CD45RA-), granulocyte monocyte progenitors (GMP; Lin-CD34+CD38+CD123+CD45RA+) and megakaryocyte erythroid progenitors (MEP; Lin-CD34+CD38+CD123-CD45RA-) (Figure 1B and 1C).

**Figure 1.**
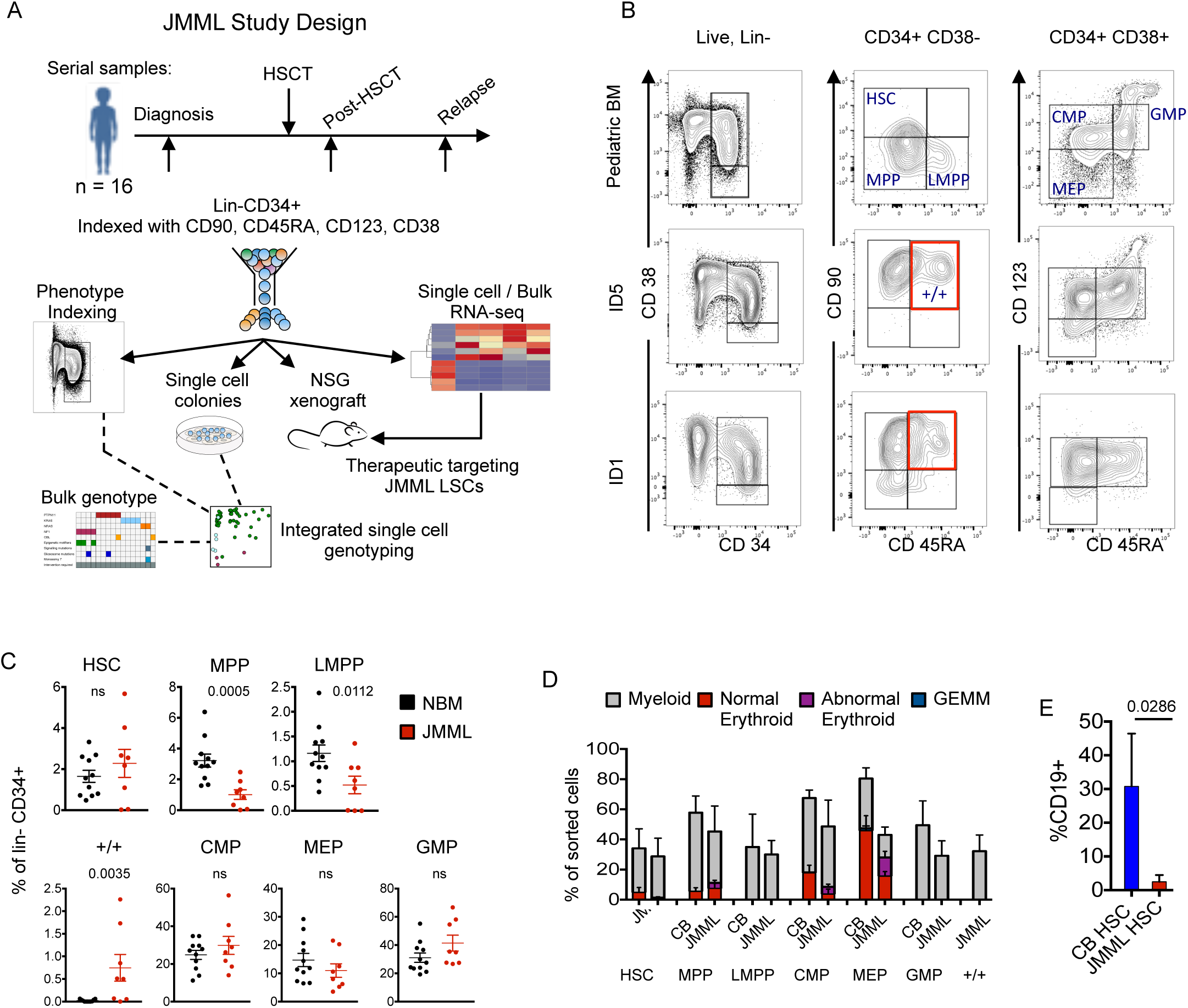
Single cell phenotypic and functional analysis reveals heterogeneity of HSPCs in JMML. (A) Outline of experimental design used in this study. (B) Representative FACS profiles of samples of normal pediatric bone marrow (BM; n=11; age 2-14 years) (top panels); BM samples from 2 JMML patients (ID5, ID1) at diagnosis (middle and lower panels). Samples gated on singlets, live and Lin-cells (left column); CD34+CD38-cells (middle); CD34+CD38+ cells (right). (C) Frequency of immunophenotypically-defined HSC and progenitors in total Lin-CD34+ cells from normal pediatric BM (n= 11; black dots) and JMML BM (n= 8; red dots); bars represent mean+SEM. (D) Clonogenic output of single HSC and progenitor cells sorted into methylcellulose; expressed as the % of sorted cells which generated a colony and the type of colony generated. Mean + SEM for 4 JMML BM samples compared to 4 normal (cord blood; CB) controls. (E) Lymphoid output of sorted HSC (100 cells) from 4 JMML BM samples (red bar) compared to 4 normal controls (CB; blue bars). Cells cultured on MS-5 stromal lines and output measured as %CD19+ cells on gated hCD45+ cells (mean+SEM) after 4 week

At a functional level, clonogenic assays of bulk BM (Figure S1A) or purified single HSPCs (Figure 1D and S1B) showed preserved myeloid output but aberrant erythroid potential of CMP and MEPs from JMML patients in comparison with cord blood (Figure 1D) and pediatric BM (Figure S1B). Erythroid colonies derived from JMML patients were frequently dysplastic (Figure S1C), in keeping with anemia seen in all patients (Table S1). The aberrant +/+ population showed exclusively myeloid output (Figures 1D and S1B). Lymphoid potential of JMML HSCs was reduced (Figure 1E) and megakaryocytic potential severely reduced (Figure S1D) in keeping with patients’ thrombocytopenia (Table S1).

Although FACS analysis supported relative preservation of a number of HSPC subpopulations in JMML, bulk phenotypic analysis may mask underlying heterogeneity of HSPCs in JMML in comparison with normal hematopoiesis. We therefore next assessed the global cellular architecture of JMML Lin-CD34+ HSPC in an unbiased manner by high throughput single-cell RNA-sequencing of 17,547 single HSPCs from JMML (n=2) and cord blood (n=2). Using 1,127 selected genes showing high level of dispersion (Figure S2A), t-distributed stochastic neighbor embedding showed highly distinct clustering of JMML HSPCs (Figure 2A) which were not equally distributed between the identified HSPC subpopulations in comparison with cord blood HSPCs (Figures 2B and S2B). JMML-specific clusters of HSPCs showed upregulation of myeloid genes (*MS4A3* and *MPO*), stem cell and fetal genes (*THY1*, *ZFP36L1, HMGA2*), proliferation markers (*MKI67*) and aberrant expression of leukemia (*HOPX*, *FOS*) and erythroid differentiation associated (*GATA1*) genes (Figure 2C, 2D). Taken together, these findings support pnenotypic evidence of HSPC compartment disruption in JMML, with aberrant myeloid bias and distinct molecular signatures.

**Figure 2.**
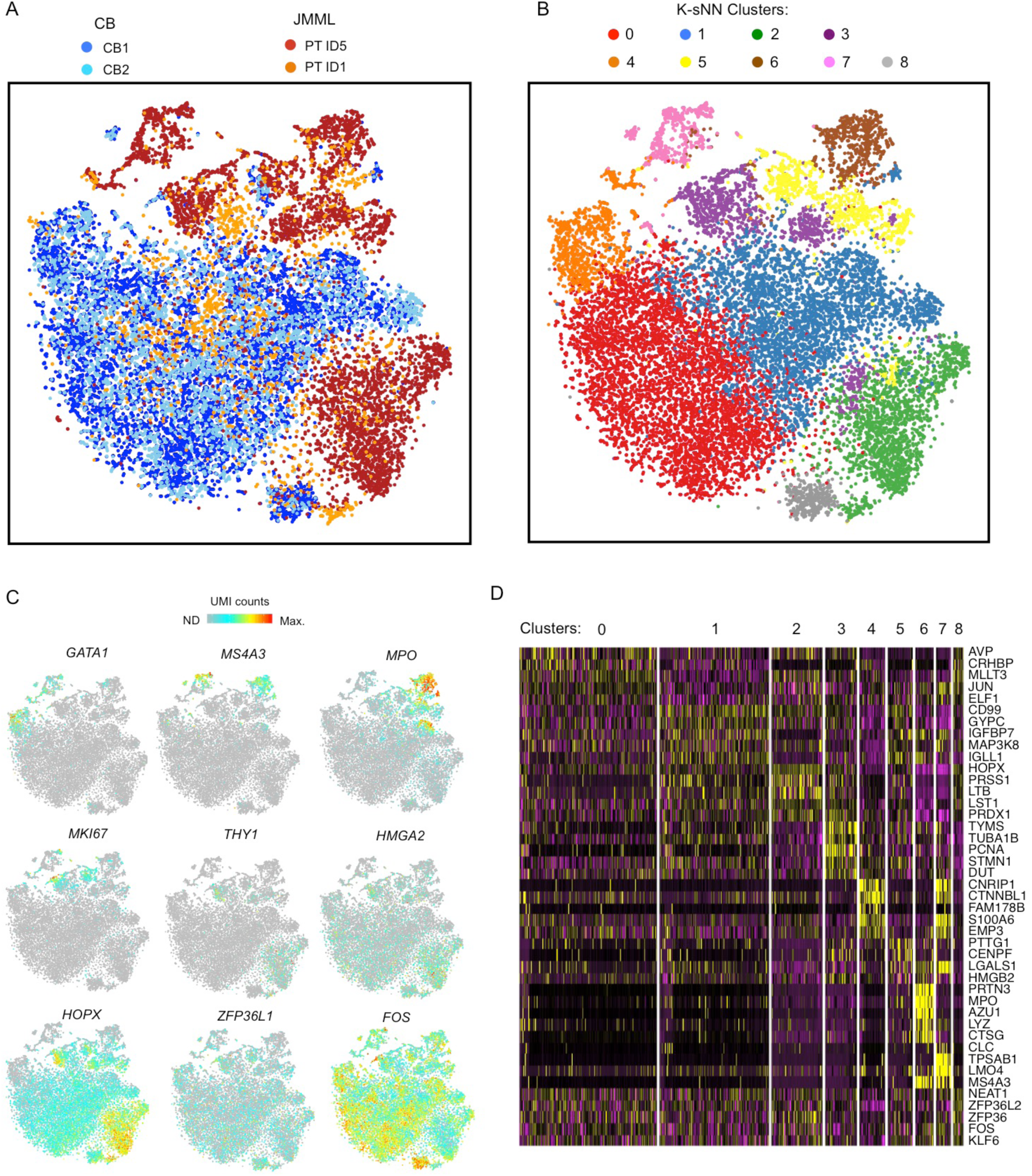
Single cell RNA-seq of JMML HSPCs reveal distinct clustering. (A) tSNE analysis of Lin-CD34+ single cell RNA-sequencing dataset based on 1,127 variable genes. 17,547 single cells that pass filtering from two CB controls (light and dark blue) and two JMML BM samples (orange and red) are shown. (B) Single cell RNA-seq clusters identified by k-shared nearest neighbor method (k-SNN) and projected by tSNE plot. Colors correspond to computationally identified clusters of single cells. (C) Selected highly variable genes from top principal components plotted on tSNE analysis from (A). Color scale represents normalized unique molecular identifier counts (UMI, grey not detected, red maximum count). (D) Heatmap depicting the top 5 positively identifying genes that distinguish each cluster of single cells shown in (B).

### Somatic mutations originate exclusively in the phenotypic HSC compartment in JMML

As driver mutations must be present in cells with self-renewal capability in order to propagate the disease, we next set out to track the cellular origin of somatic mutations within the HSPC cellular hierarchy in JMML in order to gain insights into the identity of JMML LSCs. In adult MDS, this approach has been used to identify phenotypic HSCs as the population with stem cell properties (*7*). In contrast, in AML, some mutations can be tracked to progenitor cell populations, but are absent in HSCs, supporting presence of aberrant stem cell properties in progenitor cells in AML (*10*). We first carried out a targeted mutation analysis of JMML-associated mutations in our cohort of 16 JMML cases. RAS pathway activating mutations were present in all patients while secondary spliceosome and epigenetic mutations were identified in 7/16 (44%) of patients, including all those with NF1 mutations as previously reported (*2*) (Figure 3A).

**Figure 3.**
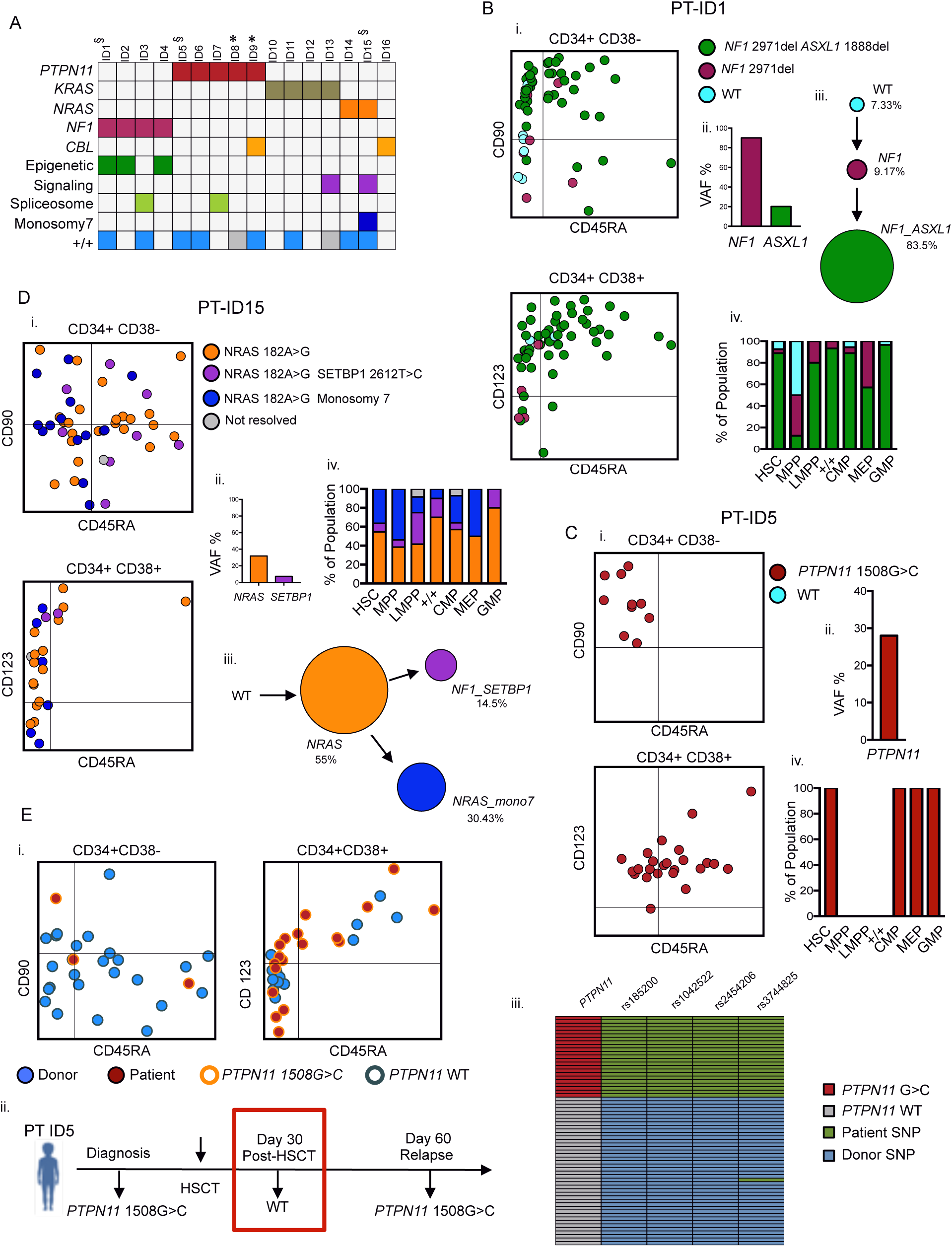
Single cell mutation tracing to phenotypic HSCs in JMML. (A) Mutations detected through a customized MDS/JMML targeted sequencing panel in whole JMML BM and/or peripheral blood samples (n=16) at diagnosis except where indicated; each column represents an individual patient. All patients had RAS gene pathway mutations; in one case involving 2 different RAS pathway genes (ID9). Secondary mutations were detected in 7/16 in epigenetic regulators (*ASXL1*,*TET2*); signaling (*SETBP1)* or spliceosome (*SRSF2*) genes (see Supplementary Table 1). Patients where the aberrant +/+ population was detected by FACS are shown in the bottom row (grey boxes = FACS not done). § samples for single cell genotyping; * relapse or early post-treatment samples. (B-D)Single cell mutational profiling for 3 JMML BM samples (ID1 (B), ID5 (C) and ID15 (D)). For each panel: i) FACS indexing of CD34+ single cells with points colored by mutational status; top CD38-, bottom CD38+; ii) VAF at diagnosis for each mutation detected in bulk BM; iii B,D) order of mutations acquired; iv) inferred mutation hierarchy and proportion of each subclone. (E) i) FACS indexing of post-transplant (day +30) BM CD34+ cells from Patient ID5 with single cells colored by donor/patient and mutational status; left CD38-, right CD38+. ii) Timeline of pre- and post-transplant *PTPN11* mutation detection by targeted sequencing of bulk BM cells showing that the mutation was not detected at day+30. Marked day 30 sample used for single cell genotyping. iii) Heatmap of the 65 single cell-derived colonies analyzed showing *PTPN11* mutational status and patient/ and donor-derived SNPs.

To track these disease-causing mutations to the HSPC hierarchy, single Lin-CD34+ cells from three patients were index-sorted into methylcellulose colony-forming assays and individual colonies picked for targeted genotyping (n=498); the index-sorting data allowed us to derive the FACS phenotype, and hence HSPC population of origin, of each colony (Figure S3A). Genotyping of SNPs demonstrated the low allelic dropout of this assay, with no false positive mutations seen in control samples (Figure S3B-H). Patient ID1 showed evidence of linear evolution with acquisition of an *NF1* followed by an *ASXL1* mutation, both of which could be tracked to all HSPC subpopulations, including HSCs (Figure 3B). The few residual wild-type cells were present in the HSC compartment and enriched in MPPs whereas mutation-positive cells were infrequent in MPPs, in keeping with phenotypic data (Fig 1C). ID5 was characterized by *PTPN11* mutation alone that was again traceable to HSCs as well as CMP and GMPs (Figure 3C). Patient ID15 showed evidence of branching clonal evolution within the HSC compartment, with a first hit NRAS mutation followed by acquisition of monosomy 7 and *SETBP1* mutation in separate subclones (Figure 3D). Finally, we analyzed a sample from patient ID5 taken post-HSCT at a time when the patient was in clinical remission but subsequently suffered overt evidence of relapse of JMML (Figure 3E). Parallel genotyping of the *PTPN11* mutation together with donor and recipient specific SNPs demonstrated presence of *PTPN11* mutation-positive HSPCs (including a single HSC) which predate (and potentially might have helped to predict) subsequent relapse. Notably, mutant positive progenitors (CD38+) were more prominent than HSCs post-HSCT, raising the possibility that these cells may predominantly drive relapse. Taken together, these data support the conclusion that all somatic mutations driving JMML disease initiation and evolution originate in the phenotypic HSC compartment with *RAS*-activating mutations as a “first hit”.

### Heterogeneity of LSCs in JMML

To further characterize JMML-LSCs, we next carried out *in vitro* and *in vivo* stem cell assays to assess self-renewal potential of JMML-HSPC subpopulations. Long-term culture-initiating cell (LTC-IC) assays showed LTC-IC potential was present in both HSCs and +/+ populations in JMML, but was absent in GMPs while in cord blood controls LTC-IC potential was restricted to HSCs, as expected (Figure 4A). We then performed xenotransplantation of purified HSCs, +/+ and GMPs, or total CD34+ cells from 4 JMML patients as shown in Figure 4B, with results of serial readout of engraftment shown in Figures 4C and D. JMML CD34+ HSPCs were highly efficient at supporting reconstitution in this NSG xenograft model, unlike adult MDS (*7*). Lineage analysis (Figure S4A) showed that transplantation from JMML donors resulted in highly myeloid-biased reconstitution in comparison with cord blood. In all cases, mice showed reconstitution following transplantation of purified HSCs from cord blood or JMML. Engraftment potential of specific JMML-HSPC subpopulations (+/+ and GMP), however, showed marked inter-patient heterogeneity of JMML-LSCs (Figure 4C and D). In patient ID1, JMML-LSC potential was restricted to the phenotypic HSC compartment. Patient ID3, who died from disease relapse following HSCT, showed similar repopulating potential in all populations (HSCs, +/+ and GMP). Patient ID5, who also relapsed after HSCT, showed most robust reconstitution with HSCs, but late reconstitution was also observed with +/+, and total CD34+CD38+ cells. It is also noteworthy that at relapse post-HSCT, the mutant positive JMML HSPC compartment in patient ID5 was dominated by mature progenitor cells (Figure 3E) unlike at diagnosis when HSCs were frequent (Figure 3C). This is consistent with relapse being primarily propagated by progenitors rather than HSCs in this patient. Finally, patient ID15 showed reconstitution following transplantation of HSC and +/+, but not GMP.

**Figure 4:**
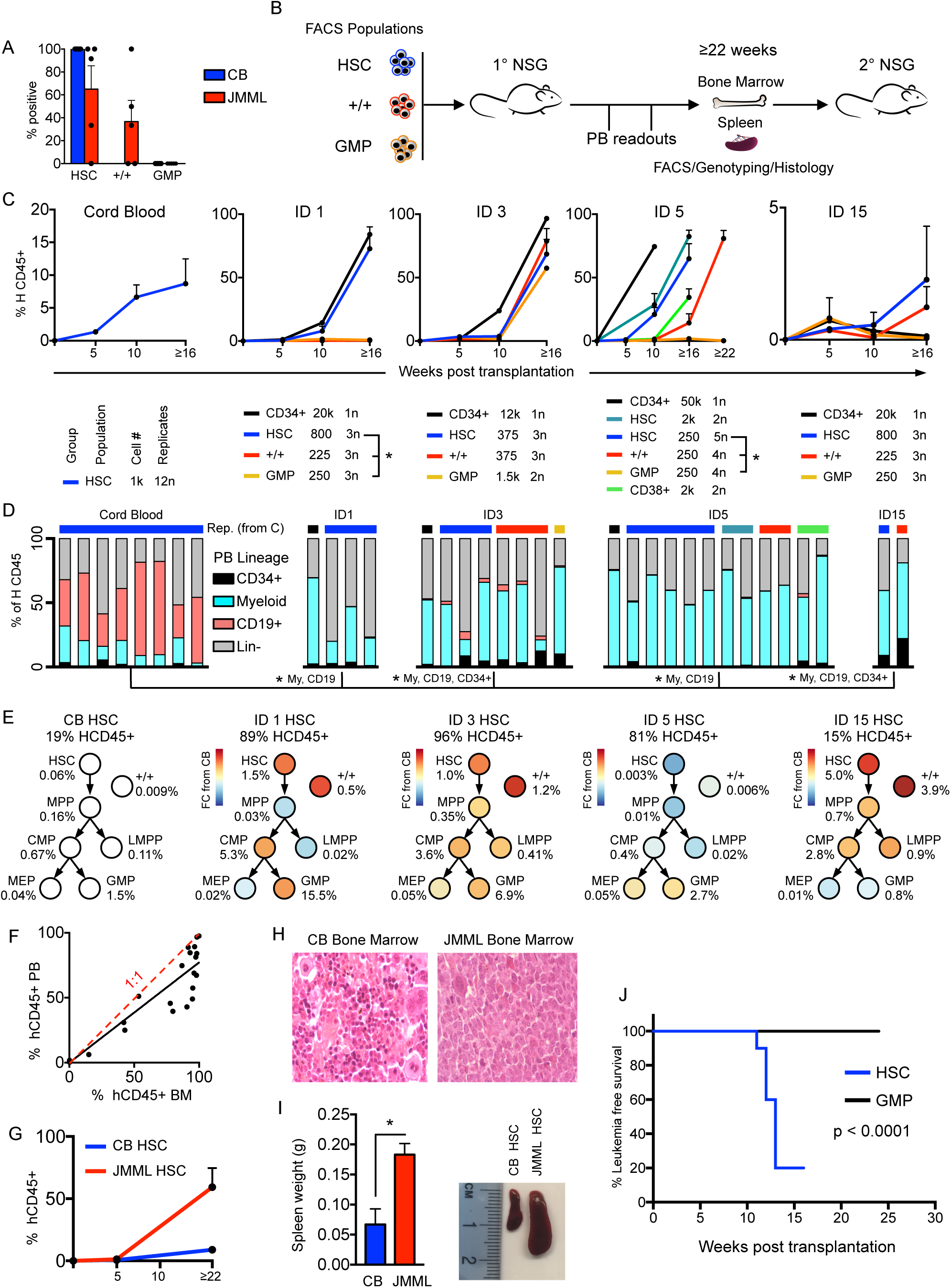
Heterogeneity of LSCs in JMML. (A) Long term culture-initiating cell (LTC-IC) assay. 100 cells from each population from 6 JMML BM samples and 5 CB controls sorted and cultured for 8 weeks. Results shown as the % of cells from each sample with clonogenic activity. (B) Schematic of *in vivo* transplantation experiments with purified HSPC populations into NSG mice. (C) Percentage of human (H)CD45+ cells in peripheral blood from xenograft CB and JMML HSPCs. Key for each patient (below) for population, number of cells transplanted, and number of replicates. *, p < 0.05 students t-test. (D) Lineage of peripheral blood reconstitution of mice with greater than 1% human engraftment at terminal time point. Color bars (top) correspond to population and cell number groups as in (C). *, p < 0.05 students t-test, compared to CB controls. (E) BM analysis of mice transplanted with HSCs from CB (n=12) or JMML BM (n=4 samples; 16 mice). Mean % human engraftment at terminal timepoint (top); % engraftment (of total HCD45+ cells) for each HSPC population shown below each population; and log fold-change for JMML compared to CB controls indicated by the color chart. (F) Comparison of % HCD45+ cells in peripheral blood versus BM at terminal time point with best fit (black) and a 1:1 ratio (red). (G) Peripheral blood HCD45+ cell % in secondary transplantation of CB and JMML HSCs. (H) Photomicrograph of Giemsa-stained BM sections from mice transplanted with purified CB (left) or JMML (right) HSC. (I) Terminal spleen weight from mice transplanted with CB vs JMML HSCs (left panel), *p < 0.05 students t-test; representative image of increased spleen size in JMML vs CB HSC-transplanted mice (right panel). (J) Kaplan Meier leukemia-free survival curve of mice transplanted with purified JMML HSCs or GMPs. Significance by log rank test.

Terminal analysis of reconstituted mice showed relative enrichment of HSCs and +/+ cells relative to cord blood engrafted animals in 3 of 4 cases analyzed (Figure 4E and Figure S4B), with higher engraftment in BM than PB (Figure 4F). JMML-HSCs also strongly supported robust reconstitution in secondary transplantations (Figure 4G). Mice transplanted with JMML-HSCs developed a JMML-like disease with cytopenias (Figure S4C), characteristic histology (Figure 4H), marked splenomegaly (Figure 4I) and reduced leukemia-free survival (Figure 4J). Taken together, these findings demonstrate that JMML-LSC activity is present in, but not confined to, the phenotypic HSC compartment. JMML-HSCs nevertheless not only showed the most robust reconstituting potential, but were also the only population able to induce JMML-like disease *in vivo* within 24 weeks and were the cell of origin of all JMML-associated driver mutations and clonal evolution events.

### Conservation of Molecular Hierarchy of JMML-HSPCs

To characterize transcriptomic signatures of JMML-LSCs we next carried out RNA-sequencing analysis of three different JMML-LSC populations (HSC, +/+ and GMP) from six JMML patients. JMML HSC showed a higher number of differentially expressed genes in comparison with cord blood HSC (n=5) than in comparison with different JMML HSPC subpopulations (Figure 5A) and, notably, +/+ cells from JMML shared molecular signatures with both HSCs and GMP (Figure S5A). Expression of known HSC genes was higher in HSCs than other populations in both JMML and cord blood, with expression in JMML +/+ cells intermediate between HSC and GMP, suggesting that a hierarchy may be preserved in JMML (Figure 5B). In keeping with this, myeloid-associated genes were more highly expressed in GMPs in both cord blood and JMML, with +/+ cells again showing intermediate expression (Figure 5C). Analysis of cell cycle genes supported that quiescence-associated gene expression was higher in cord blood HSCs with proliferation-associated transcriptional changes in all JMML HSPC populations, including HSCs, but more so in JMML GMPs (Figure 5D). Interestingly, the fetal HSC genes *HMGA2*, *CNN3*, and *VNN2* were highly expressed in JMML HSCs and to a lesser extent in JMML +/+ and GMP cells (Figure 5E). Gene set enrichment analysis (GSEA) further supported that JMML-HSCs showed more HSC- and quiescence-associated gene expression than JMML-GMPs (Figure 5F). In contrast, JMML-GMPs showed upregulation of proliferation and pediatric cancer signatures in comparison with JMML-HSCs (Figure 5F). Direct comparison of JMML versus cord blood HSCs showed a number of clusters of aberrantly expressed genes, clearly distinguishing JMML HSCs (Figure 5G). Hallmark GSEA revealed enrichment of a number of proliferation-associated (G2M checkpoint, MYC and E2F) and DNA repair gene sets in JMML-HSCs versus cord blood HSCs (Figure 5H). We identified a core set of 24 genes which distinguished JMML-HSCs from JMML-GMPs and cord blood HSCs (Figure 5I) and were overexpressed in JMML-specific HSPC subpopulations identified by single cell RNA-sequencing (Figure S5B). This set of genes included the non-DNA-binding homeodomain protein HOPX, a regulator of primitive hematopoiesis (*11*) and the serine/threonine kinase STK24, a target of the kinase inhibitor bosutinib (*12*) (Figure 5J).

**Figure 5.**
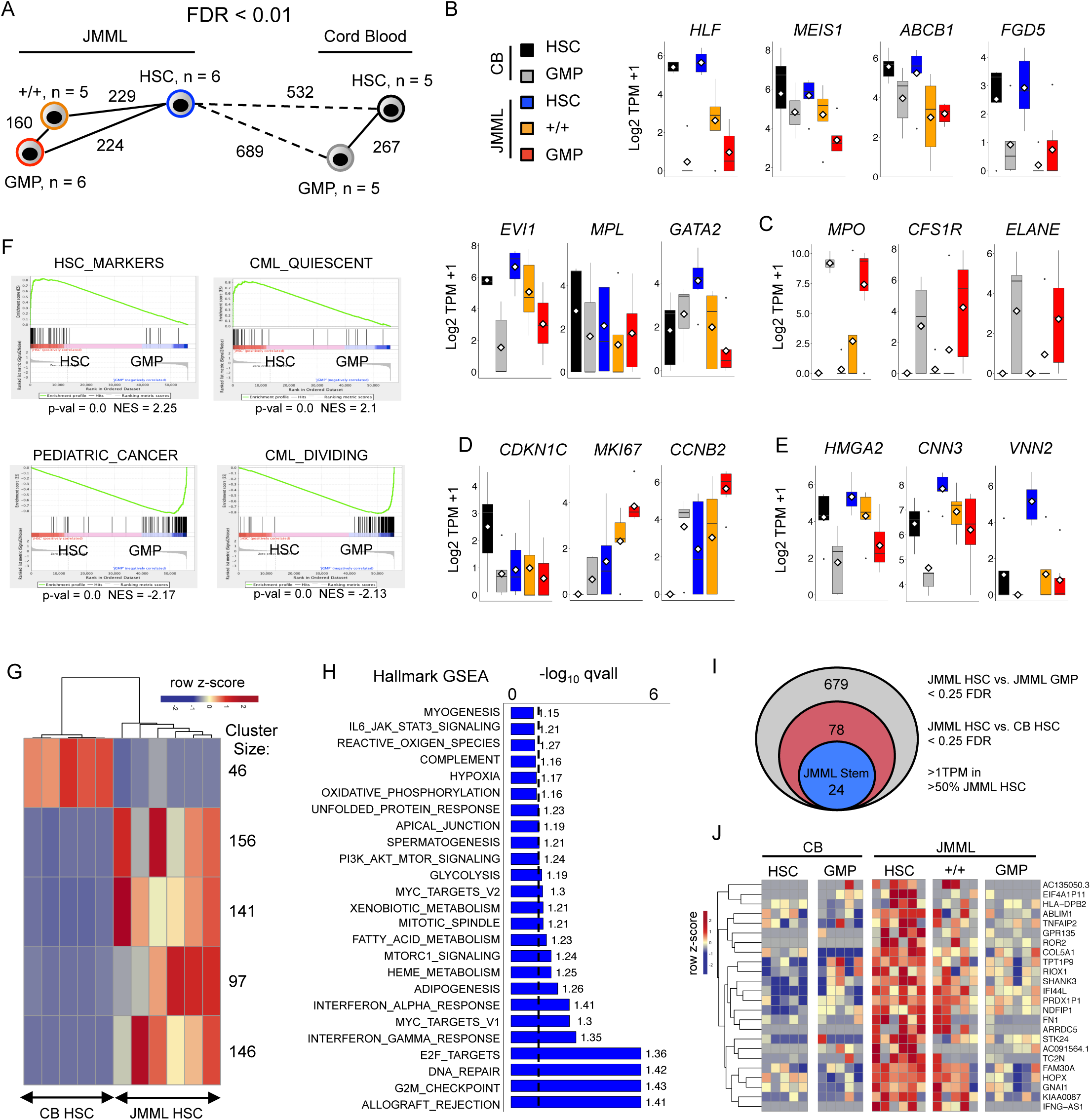
Conservation of Molecular Hierarchy of JMML-HSPCs. (A) Nearest neighbor analysis of purified HSPC populations from CB and JMML. Numbers represent significant differentially expressed genes <0.1 FDR. (B-E) Gene expression for CB HSCs (black), CB GMPs (grey), JMML HSCs (blue), JMML CD90+/CD45RA+ (+/+; orange), and JMML GMPs (red) for known stem cell genes (B); myeloid genes upregulated in JMML and CB GMPs (C); cell cycle-related genes (D); expression in JMML HSPCs; known fetal HSC genes (E). (F) Gene set enrichment analysis of JMML HSCs vs JMML GMPs with upregulation of gene sets for HSC markers and CML quiescence in HSCs (top), and upregulation of gene sets for pediatric cancer and CML proliferation gene sets in GMPs (bottom). NES; Normalized enrichment score. (G) Heatmap depicting k-means clustering of top differentially expressed genes between JMML HSCs and CB HSCs with an FDR cutoff < 0.1. Cluster size corresponds to the number of genes within each significant cluster. (H) Hallmark GSEA Between JMML HSCs and CB HSCs depicting upregulated gene sets in JMML. Dotted line represents an FDR cutoff of < 0.1; numbers next to the bars represent NES. (I) Identification of a JMML LSC specific gene signature from bulk RNA-sequencing. (J) Heatmap showing expression of the 24-gene JMML LSC signature from (I) in CB and JMML HSCs, GMPs and +/+ cells

### Novel Therapeutic Targets for JMML-HSCs

A number of putative therapeutic targets were upregulated in JMML-HSCs, including overexpression of *SLC2A1* (GLUT1) and RAS-associated transcription (Figures 6A and B). Specific targeting of GLUT1 with Fasentin or of RAS-associated signaling with the MEK inhibitor PD901 both significantly and differentially (in comparison with cord blood) reduced clonogenicity of JMML HSCs (Figure 6C). Bromodomain inhibitors have been shown to reverse RAS-associated transcriptional changes (*13, 14*), and we also demonstrated significant differential inhibition of JMML-HSC clonogenicity with JQ1 (Figure 6C).

**Figure 6.**
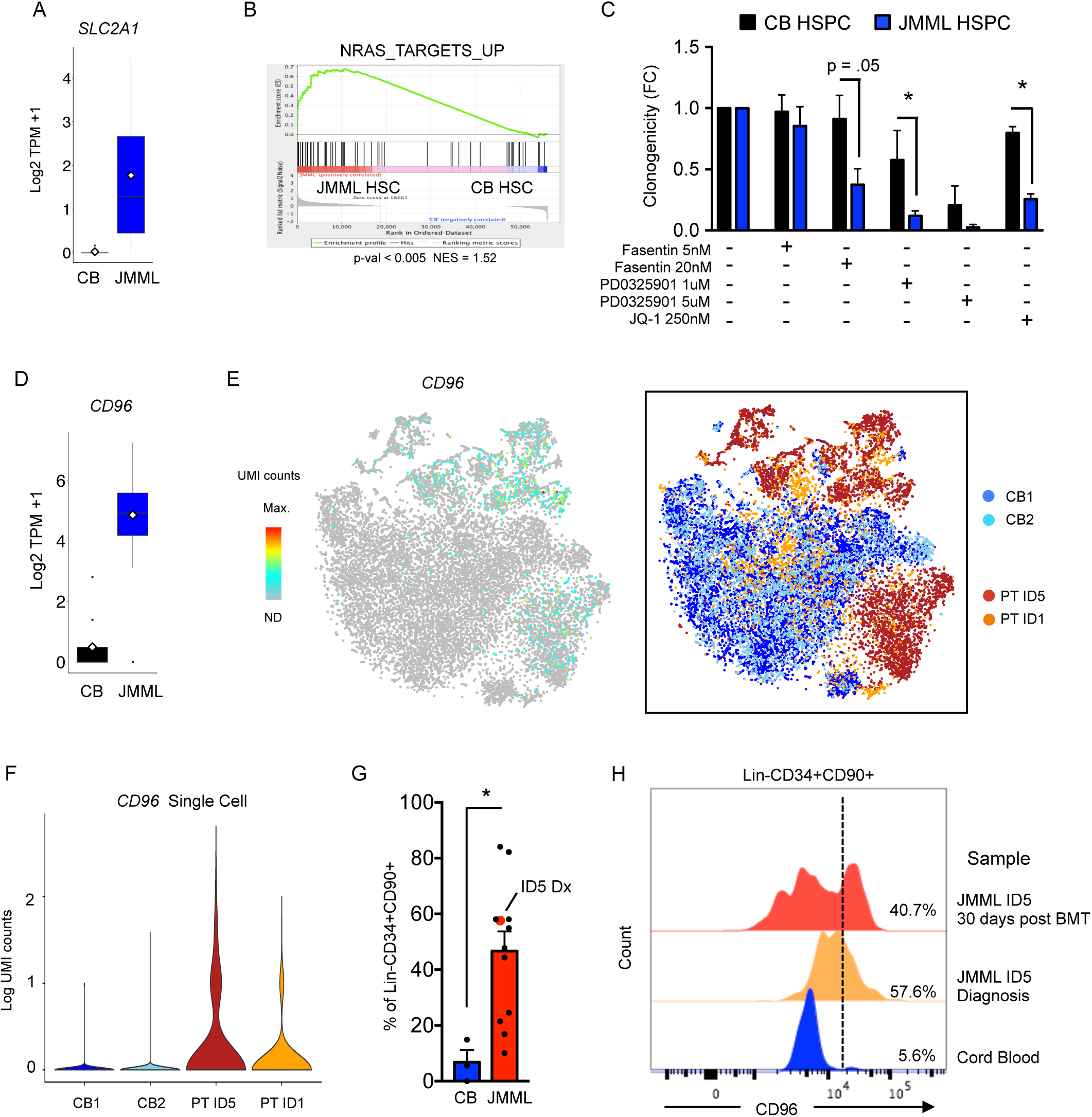
Novel Therapeutic Targets for JMML-HSCs. (A) Gene expression of selected potential therapeutic targets significantly upregulated in JMML HSCs vs CB HSCs. p < 0.05 Benjamini-Hochberg corrected FDR. (B) Gene set enrichment analysis for NRAS targets between JMML HSCs and CB HSCs. *, p < 0.05 students t-test. (C) In vitro clonogenicity of CB and JMML CD34+CD38-HSPCs with selected targeted inhibitors. (D) CD96 expression by bulk RNA seq in JMML and CB HSCs. (E) CD96 expression by single cell RNA-seq (left) on displayed on tSNE analysis from CB and JMML (right). (F) Violin plot of single cell CD96 expression in JMML and CB CD34+ HSPCs. (G) CD96 expression by FACS in CB and JMML CD34+CD90+ cells showing % of LIn-CD34+90+ cells expressing surface CD96+. (H) Presence of CD96+ cells in paired diagnostic and post-transplant samples in Patient ID5

One of the top differentially expressed genes (overexpressed in JMML-HSCs) was the cell surface receptor CD96 (Figure 6D), also confirmed to be more frequently expressed in JMML-specific subpopulations of HSPCs in our single cell RNA-seq dataset (Figures 6E and F). CD96 has previously been proposed as a biomarker and therapeutic target in AML (*15*). We confirmed aberrant surface expression of CD96 protein on JMML-LSCs (Figure 6G) and show that this aberrant population of CD96-expressing cells could be detected in a patient post-HSCT who subsequently relapsed (Figure 6H), raising the possibility that CD96 could be a useful biomarker for JMML-LSCs.

Taken together, these data support that JMML-LSCs are heterogeneous, with multiple different HSPC populations showing capacity for self-renewal. JMML driver mutations originate in HSC-like cells which reside at the apex of the JMML-LSC hierarchy, and this population shows increased reconstitution potential and HSC-associated gene expression in comparison with other JMML-HSPC populations.

## Discussion

Disease relapse after achievement of clinical/morphological remission is a major cause of treatment failure across many different human cancers. Characterization of distinct CSCs, specific to each type of malignancy, is a crucial step towards improved approaches for disease monitoring in order to predict disease relapse and the development of CSC-directed therapy. This task is difficult in rare pediatric diseases such as JMML, and yet the need for new disease biomarkers in this disease is particularly acute because HSCT, the only curative therapy, carries a high rate of relapse that is often difficult to diagnose promptly. In hematopoietic cancers such as MDS, MPN, ALL and AML (*4, 5, 7-9*), LSCs are diverse and vary considerably in their frequency and phenotype, with each providing important insights into LSC biology. However, in the rare and poor prognosis childhood leukemia JMML, very little is known about the cellular hierarchy and identity of LSCs, including whether progenitor cells are transformed in this condition. We have carried out a comprehensive phenotypic, functional and molecular analysis of a cohort of JMML patients and describe significant disruption of the HSPC hierarchy in JMML, including presence of an aberrant early progenitor cell (Lin-CD34+CD38-CD90+CD45RA+) which may reflect an underlying aberrant HSC myeloid differentiation pathway in JMML. Interestingly, this population has also been observed in adult patients with myeloid malignancies associated with monosomy 7 (*16*). We also observed a reduction of LMPPs, a key population of early lymphoid progenitors, as well as reduced lymphoid potential of HSCs, supporting that the myeloid phenotype associated with JMML may be driven by a myeloid-bias already present in the HSC. Consistent with this, we also observed impaired megakaryocyte and erythroid output from JMML HSC.

Backtracking of somatic genetic lesions to distinct HSPC subsets is a powerful method to help identify CSCs, as mutations acquired by short-lived progenitors which lack self renewal ability are not able to propagate the disease (*7*). Using a similar approach, combining single-cell index sorting and colony genotyping, we demonstrate that all JMML driver mutations could be backtracked to the phenotypic HSC compartment with *RAS*-activating mutations as a “first hit”. Despite being relatively genetically simple in comparison with other tumors, we observed JMML patients with both linear and branching patterns of clonal evolution originating in JMML-HSCs. Importantly, we were able to detect mutant HSPCs in a post-HSCT patient a month before molecular/clinical evidence of relapse. At the same time point, a population of CD96+ cells within the Lin-CD34+CD90+ gate could be detected by FACS on patient cells but not normal controls. Taken together, this strongly suggests that the presence of JMML-HSPCs with aberrant phenotype could be used to predict impending relapse in JMML patients, thus providing a much needed biomarker of residual disease post-HSCT, although this will need to be validated in larger clinical cohorts before this could be applied clinically.

Although our colony genotyping strongly supported that HSC are the cell of origin in JMML, functional analysis (xenotransplantation and LTC-IC assays) also revealed marked inter-patient heterogeneity of JMML-LSCs. Thus, while JMML HSCs were consistently able to support LTC-IC potential and engraftment in xenograft models, some patients also showed evidence of self-renewal of progenitor cells, including GMPs and the novel +/+ cell population that we describe. Interestingly, this suggests that JMML displays a distinct biology, sharing some features with MDS/MPN, which are propagated by the counterparts of HSCs (*5, 7*), and other features with acute leukemias, which show transformation of progenitor populations, but cannot usually be propagated by HSCs lacking the late driver mutations acquired by transformed progenitor/precursors (*8-10*). Furthermore, JMML-HSPC engrafted robustly in NSG mice, rather than exhibiting the poor *in vitro* and *in vivo* proliferative capacity, characteristic of adult MDS stem cells (*7*). These biological features of pediatric versus adult MDS may reflect the impact of *RAS* driver mutations on JMML-HSC and/or co-expression of a proliferative fetal gene program, such as we observed here in the JMML-HSC but not reported in adult MDS (*7*). Of specific note, the fetal HSC specific gene *VNN2*, encoding the glycophosphatidylinositol-anchored surface protein GPI-80, was markedly overexpressed by JMML-HSCs (*17*). This is of particular interest as disease-associated mutations in JMML can often be tracked back to neonatal bloodspots and fetal hemoglobin is often increased in JMML cases (*1*), together suggesting a fetal origin of JMML. However, a limitation of this current analysis is that earlier neonatal samples taken at birth (cord blood or neonatal blood spots) were not available. Consequently, whilst our data support a fetal origin of the JMML cases studied, this can not be definitively established.

Although phenotypic analysis of HPSCs showed considerable overlap with normal pediatric BM HSPCs, such analyses, based on a small number of canonical surface markers, often fail to reveal underlying cellular heterogeneity. Single cell RNA-sequencing analysis is a powerful method to resolve such heterogeneity and has been widely used to analyze normal HSPCs, but less so malignant hematopoiesis (*18*). We have recently shown that in chronic myeloid leukemia, such an approach can help to resolve normal and leukemic stem cells (*19*). We therefore used a similar approach to analyze over 17,000 HSPCs from JMML patients and cord blood, revealing that JMML-HSPC are molecularly highly distinct. Bulk RNA-sequencing analysis of JMML-LSCs revealed evidence of a hierarchical organization of HSPCs in JMML, with HSCs residing at the apex of this hierarchy and +/+ cells sharing features of both HSCs and GMPs. Importantly, this also allowed us to identify a number of upregulated or aberrantly expressed putative therapeutic targets in the JMML-HSC, including *SLC2A1* (GLUT1) and RAS-associated pathways. Furthermore, the clonogenicity of JMML-HSC was specifically reduced by targeting either GLUT1 (with Fasentin) or RAS-associated pathways (with the MEK inhibitor PD901 or the bromodomain inhibitor JQ1). As eradication of CSC/LSC is not only necessary but also potentially sufficient to achieve disease eradication (*4*), further preclinical evaluation of these targets is warranted.

In summary, we describe an integrated phenotypic, functional and molecular analysis of JMML-LSCs, illustrating marked intra- and inter-patient heterogeneity of LSCs. We identified a number of candidate biomarkers and therapeutic targets, paving the way for LSC-directed disease monitoring and therapy in JMML.

## Methods

### Patient Sample Collection

Patient samples and normal controls were prospectively collected in accordance with the Declaration of Helsinki for sample collection and use in research under the UK NIHR Paediatric MDS/JMML study (NHS REC reference number 15/LO/0961) or normal pediatric bone marrow (NHS REC reference number 12/LO/0426). An additional JMML patient sample was provided by the Bloodwise Childhood Leukemia Cell Bank (NHS REC reference number 16/SW/0219). Cord blood samples were commercially sourced (Zenbio US, Cat# SER-CD34-F). Detailed phenotypic and clinical characteristics (Table S1) were captured on a secure online platform, using the Human Ontology Database tool. Clinical details for the sample provided by the Leukemia Cell bank were not available. Mononuclear cells (MNC) from peripheral blood and BM samples were isolated on Ficoll density gradients and cryopreserved in 90% fetal bovine serum (FBS) and 10% dimethylsulfoxide (DMSO). Cells were thawed and processed for downstream analysis as previously described (Woll et al., 2014). Additional information on FACS sorting of samples are detailed in Supplementary Methods.

### In Vitro Assays

Materials and methods for *in vitro* assays including cell clonogenic assays, Erythroid / megakaryocytic culture assay, B-cell differentiation MS5 co-cultures, Long-Term Culture-Initiating Cell assay, and Mutational analysis of single cell colonies are detailed in the Supplementary Methods.

### NSG Mouse Studies

All procedures involving animals were approved by the UK Home Office Project license (PPL 30 – 3103). NSG mice were obtained through Jackson Laboratory and maintained in individually ventilated cages in a specific pathogen free facility. Male and female mice were randomly assigned to groups. NSG xenografts were carried out in 10-14 week old mice. Additional methods on analysis of NSG experiments are detailed in Supplementary Methods.

### Single Cell RNA-sequencing

18,333 – 24,000 single cells from the Lin-CD34+ population from two JMML patient samples (Patient ID1 and ID5) and 2 normal cord blood controls, were sorted and processed for RNA seq using the 10X chromium platform as per the manufacturer’s instructions (10X Genomics, Cat# 120237). In short, cells were sorted into a total volume of 40 µL and 33.8 µL of the sample was loaded onto the Single Cell 3’ Chromium 10X Chip with 66.2 µL of the 10X master mix. cDNA was generated on the Chromium 10X controller. Post RT clean up was performed as per the manufacturer’s instructions and the product was amplified with 8 PCR cycles. Post amplification clean up and library construction was performed as per the manufacturer’s instructions and samples were sequenced on the Illumina HiSeq4000, with read1 being 26bp and read2 98bp. Additional methods for analysis are detailed in Supplementary Methods.

### Bulk RNA-sequencing

FACS purified populations were processed for RNA-seq using the Smart Seq2 protocol as previously described (*20*). Briefly, fifty purified cells were sorted directly into 4 µl of lysis buffer containing 0.4% Triton X-100 (Sigma-Aldrich), RNase inhibitor (Clontech), 2.5 mM dNTPs (Thermo Fisher) and 2.5 µM oligo-dT30VN primer (Biomers.net). cDNA was generated using SuperScript II (Invitrogen), pre-amplified using KAPA HiFi HotStart ReadyMix (KAPA Biosystems) using 19 cycles of amplification. After PCR amplification, the cDNA libraries were purified with AMPure XP beads (Beckman Coulter) using a ratio of 0.8:1 beads to cDNA, according to the manufacturer’s instructions. Post purification libraries were resuspended in EB buffer (Qiagen). The quality of cDNA traces was assessed by using a High Sensitivity DNA Kit in a Bioanalyzer instrument (Agilent Technologies). Tagmentation and library preparation was performed using the Nextera XT DNA Library Preparation Kit (Illumina, Cat# FC-131) according to the manufacturer’s instructions. Samples were sequenced using the Illumina NextSeq 500 platform, generating 75 bp single-end reads. Additional information on analysis of bulk RNA-seq are detailed in the Supplemantary Methods.

## Acknowledgments

The authors thank all patients and their families for contributing samples for this research and the BMS, Oxford University, for technical assistance. This work was funded by a Medical Research Council Senior Clinical Fellowship (MR/L006340/1) to AJM, a National Institute for Health Research Fellowship to EL, a MRC-funded Oxford Consortium for Single-cell Biology (MR/M00919X/1) and the Oxford NIHR Biomedical Centre based at Oxford University Hospitals NHS Trust (IR) and University of Oxford. AR is supported by a Bloodwise Clinician Scientist Fellowship (17001), NF by a Kay Kendall Leukaemia Fund Fellowship (KKL1124) and DI by a Bloodwise Clinical Research Fellowship. The views expressed are those of the author(s) and not necessarily those of the NHS, the NIHR or the Department of Health. The authors acknowledge the contributions of the WIMM Flow Cytometry Facility, supported by the MRC HIU; MRC MHU (MC_UU_12009); NIHR Oxford BRC and John Fell Fund (131/030 and 101/517), the EPA fund (CF182 and CF170) and by the WIMM Strategic Alliance awards G0902418 and MC_UU_12025.

## Author Contributions

EL and BP performed experiments, analyzed the data and helped to write the manuscript. ARM performed experiments and analyzed data. AH and AR analyzed and interpreted data. GB, NA, CAGB, NE, DI, NS, NF and SOB conducted experiments and analyzed data. JDLF provided patient samples. AR conducted the clinical study, helped advise on experimental design and provided patient samples. IR and AJM conceived, designed and supervised the research, analyzed the data and wrote the manuscript.

---

**Figure S1.**
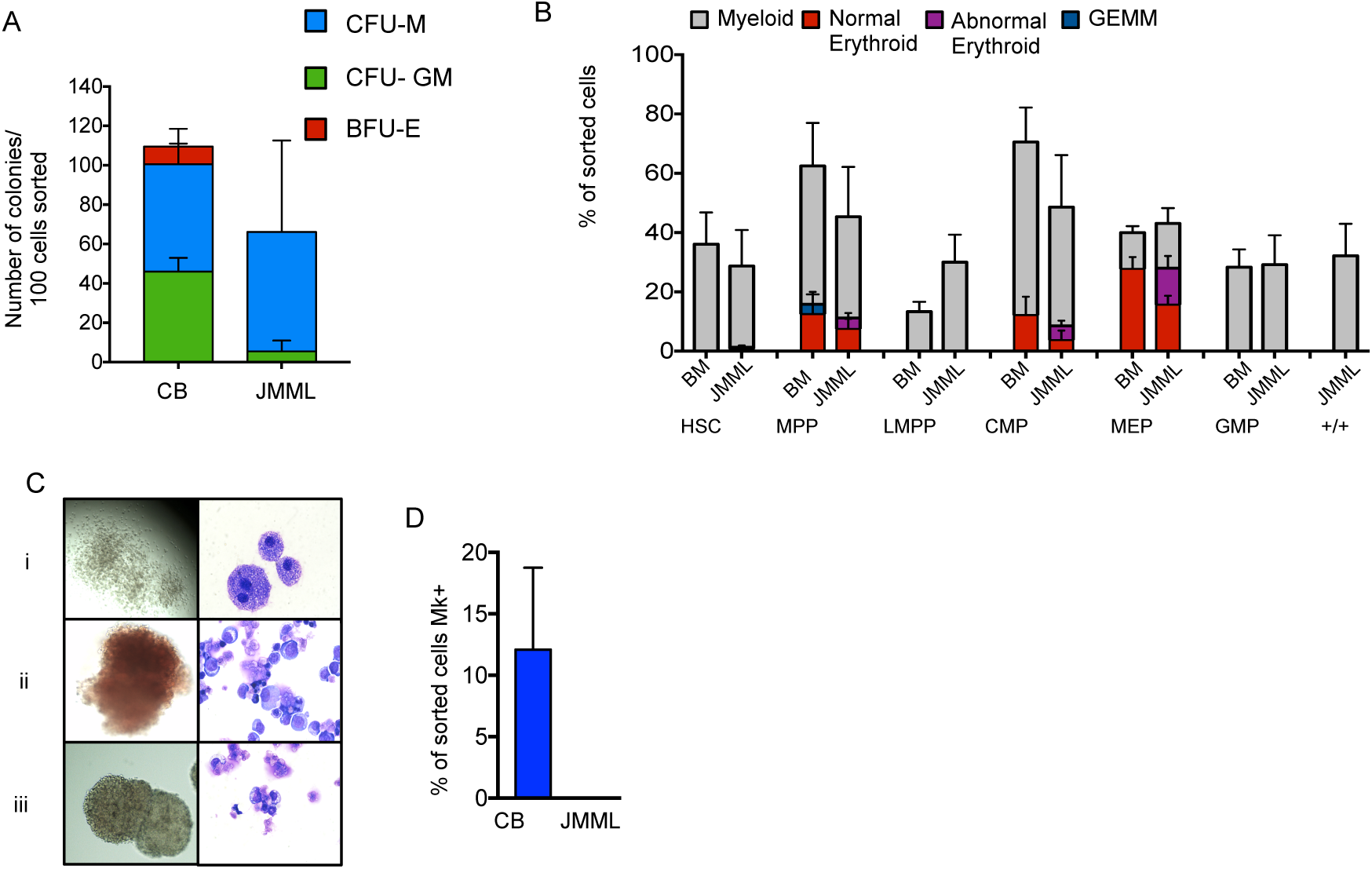
Single cell phenotypic and functional analysis reveals heterogeneity of HSPCs in JMML. (A) Bulk clonogenicity in methylcellulose, 100 cells sorted per assay (CB control n=2, JMML n=2). (B) Clonogenic output of purified HSC and progenitor populations from normal pediatric BM (n=3) and JMML (n=4) in single-cell methylcellulose assays. (C) Representative examples of colonies (left column) and MGG-stained cells from individual plucked colonies (right column) from the clonogenic assays in (B): i. macrophage colonies from the JMML Lin-CD34+CD38-CD90+CD45RA+ population ii. ‘normal appearing erythroid colonies and iii. abnormal erythroid colonies derived from JMML MEP. (D) % of wells with CD42+ megakaryocytes in single cell liquid cultures of normal CB (n=2) and JMML (n=2).

**Figure S2.**
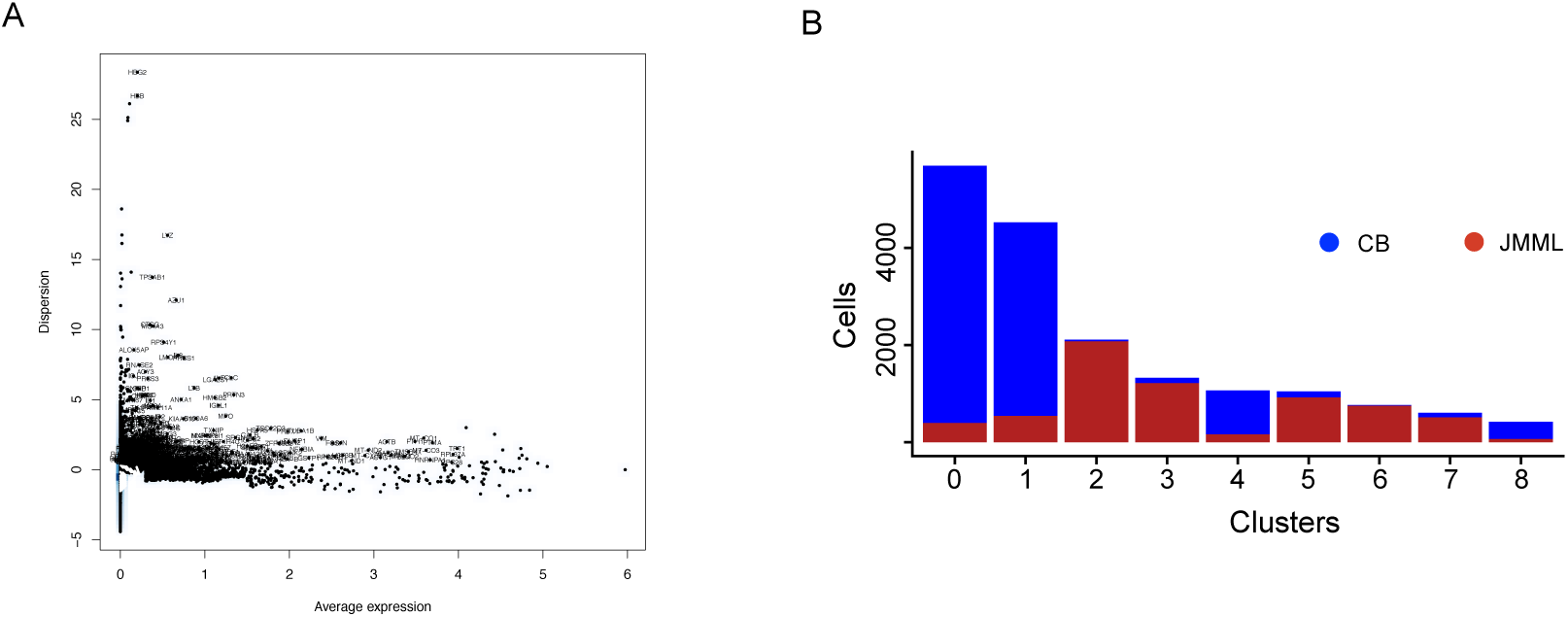
Related to Figure 2. Single cell RNA-seq of JMML HSPCs reveal distinct clustering. (A) Plot showing single cell mean gene expression versus Dispersion. Genes selected for tSNE and clustering analysis are highlighted. (B) Number of single cells within each cluster from CB (blue) and JMML (red).

**Figure S3.**
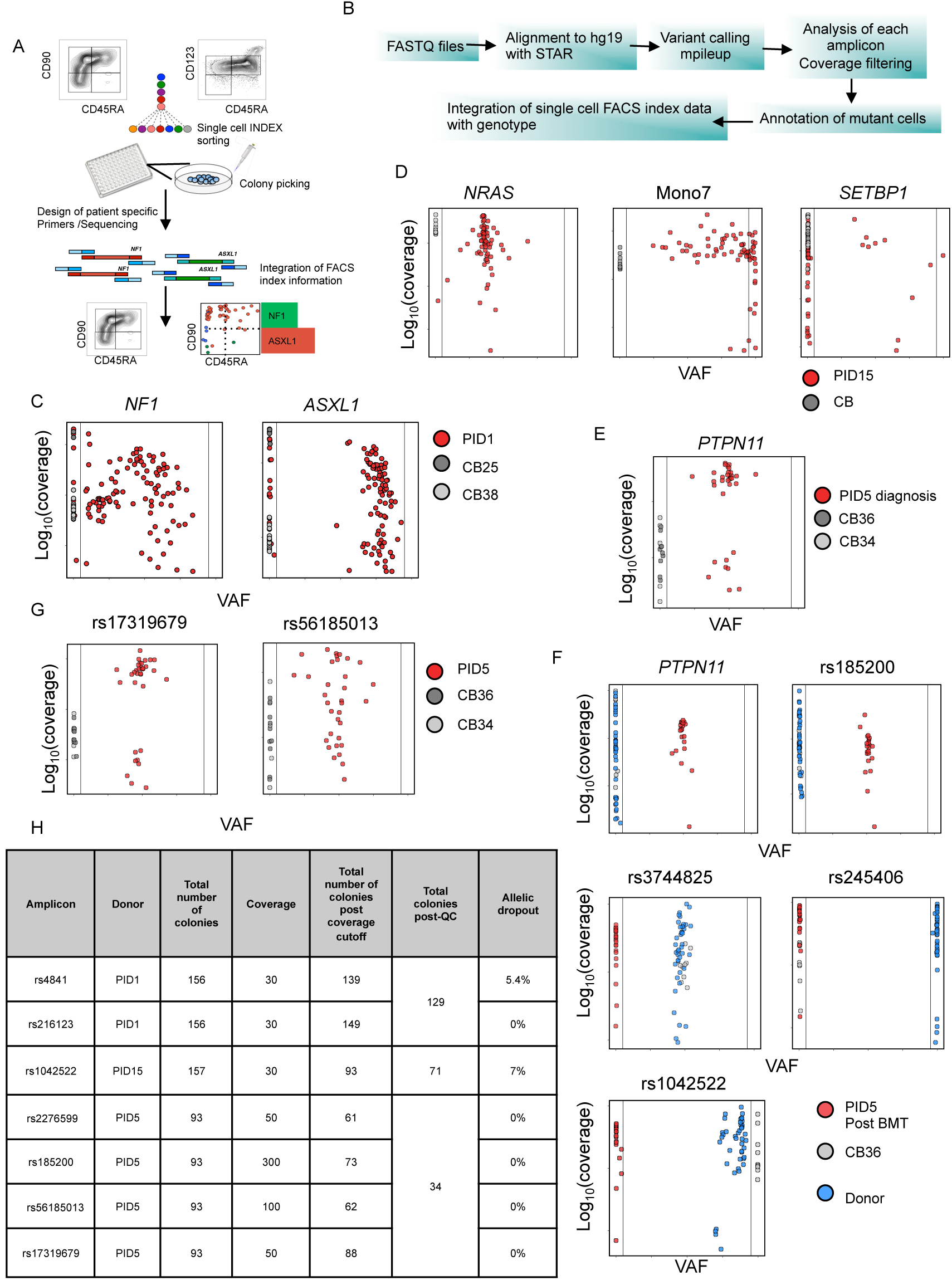
Related to Figure 3. Single cell mutation tracing to phenotypic HSCs in JMML – quality control. (A) Diagram of the workflow for single cell genotyping (B) Pathway for samples analysis (C – F) Variant allele frequency and coverage of amplicons sequenced from patient derived colonies and cord blood controls. (F) PID 1 n= 111 cells used for analysis, PID15 n=71 cells used for analysis, PID 5 n= 34 cells used for analysis, PID5 post BMT n=65 cells used for analysis (G) Examples of coverage for amplicons from heterozygous SNPs used to calculate the allelic drop out of the method (H) Summary of amplicons heterozygous SNPs for each sequencing run; coverage applied and allelic drop out for each amplicon

**Figure S4.**
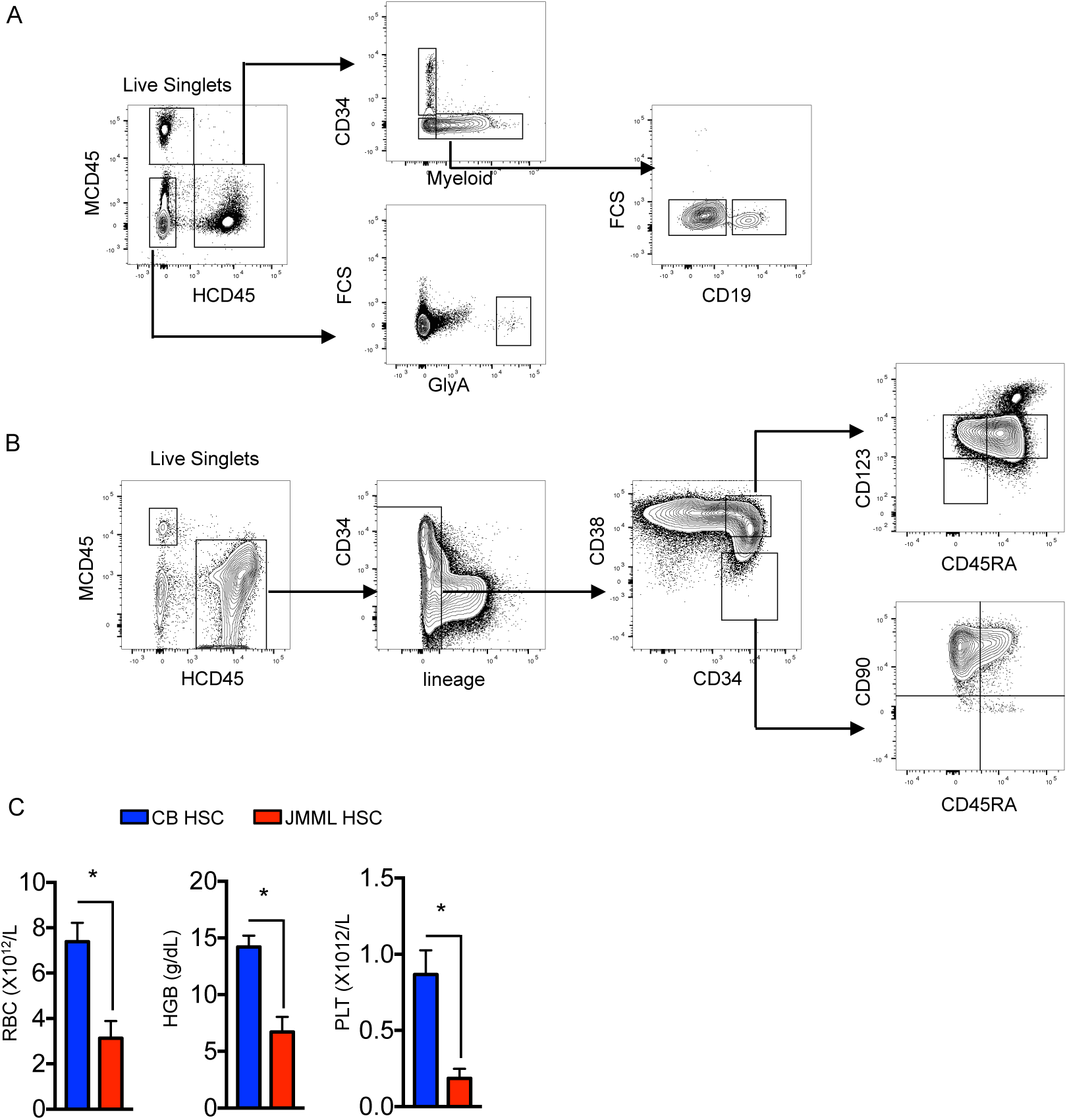
Related to Figure 4. Functional characterization of JMML HSPCs. (A) Representative FACS analysis of peripheral blood from JMML xenograft showing gating strategy. (B) Representative FACS analysis of bone marrow from JMML xenograft overall showing gating strategy. (C) Terminal peripheral blood analysis from mice transplanted with CB or JMML HSC; * p < 0.05 students t-test.

**Figure S5.**
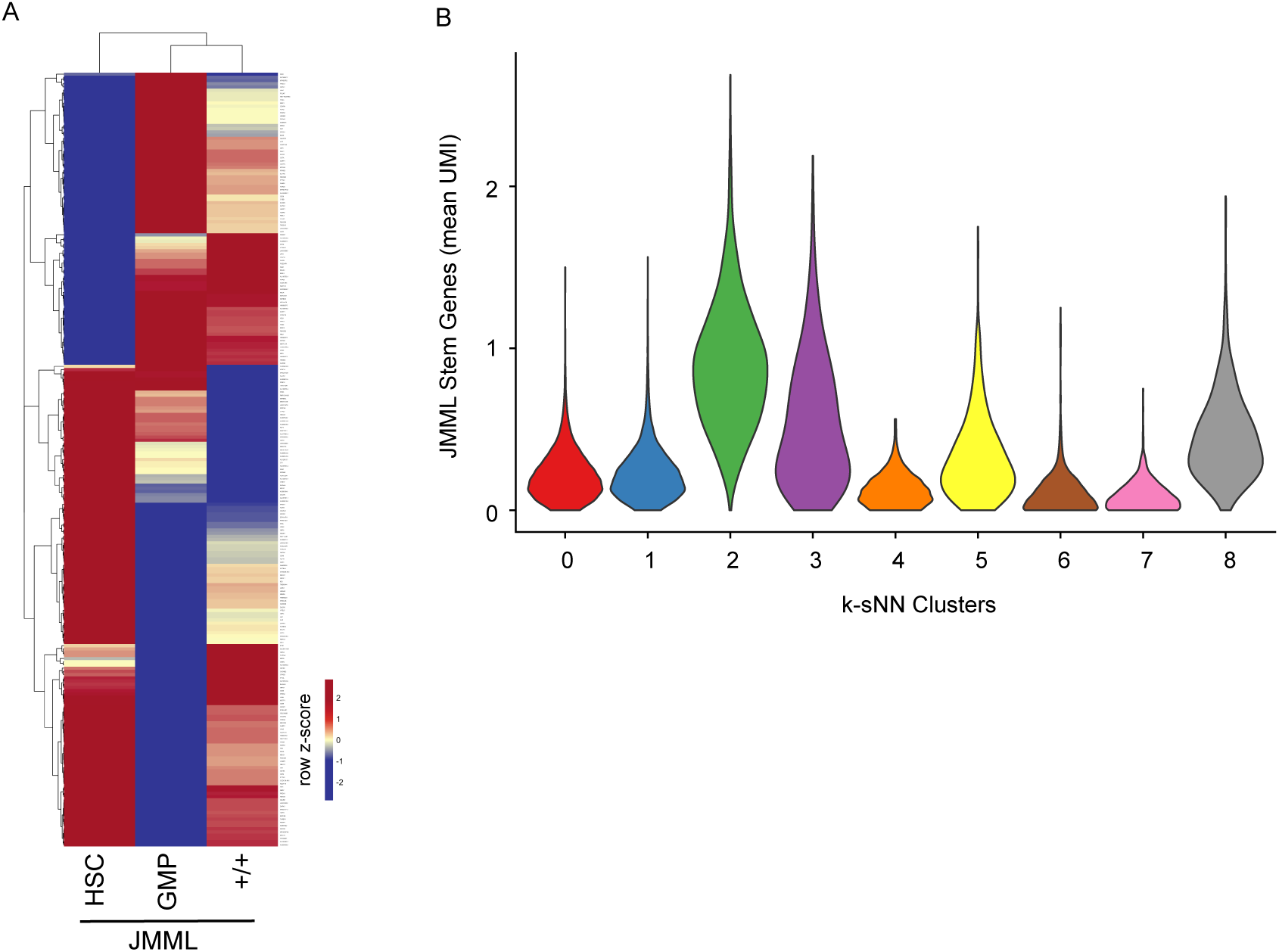
Related to Figure 5. Molecular Characterization of JMML HSPCs. (A) Heatmap of significant differentially expressed genes in JMML bulk populations. (B) Single cell gene expression of the JMML stem cell genes identified in Figure 5I. Clusters are as identified in Figure 2B. Expression plotted as mean UMI.

## Supplementary Methods

### Contact for reagent and resources

Further information and requests for resources and reagents should be directed to and will be fulfilled by Adam Mead.

### Cell lines

The M210B4 cell line was maintained in RPMI (Gibco) 10% FBS (HyClone), the Sl/Sl cell line was maintained in DMEM (Gibco) 15% FBS (HyClone), and the MS5 cell line was maintained in AlphaMEM (Invitrogen) 10% FBS (HyClone).

### Fluorescence activated cell sorting

FACS-sorting of MNCs was performed using the FACS Aria II, FACS Aria III and FACS Aria Fusion (Becton Dickinson). All FACS experiments included single color-stained-controls (CompBeads, BD Biosciences) and Fluorescence Minus One controls (FMO). Bulk and single cells HSPCs were isolated from JMML patient samples and normal controls. For all single cell experiments, sorting was performed using ‘single cell’ purity setting, tested by the deposition of single beads into the bottom of 96 well plates and checking by fluorescence microscopy. Index sort data for single cells of the mean fluorescence intensities (MFI) of CD34, CD38, CD90, CD45RA, CD123 were also recorded for each individual cell isolated. In order to check the correct alignment of the sorter, BD FACS Accudrop Beads (BD Biosciences) were sorted into a 96 well plate or eppendorf tube in order to ensure that all beads were delivered to the bottom of the well/tube. Purity of sorted population was performed by reanalysis of sorted populations and was typically >95%. Flow cytometry profiles of the human HSPC compartments were analyzed using FlowJo software (version 10.1). Antibodies for HSPC sorting and analysis were CD34 APC-eFlour780 (eBiosciences, Clone: 4H11, Cat#: 47-0349-42, RRID: AB_2573956), CD38 PE-Texas Red (Invitrogen, Clone: HIT2, Cat#: MHCD3817 RRID: AB_10392545), CD90 BV421 (Biolegend, Clone: 5 E 10, Cat#: 328122, RRID: AB_2561420), CD45RA PE (eBiosciences, Clone: HI100, Cat#: 12-0458-41 RRID: AB_10717397), CD123 PE Cy7 (Biolegend, Clone: 6H6, Cat#: 306010, RRID: AB_493576), and CD8 (Biolegend, Clone: RPA-T8, Cat#: 301006, RRID: AB_314124), CD20 (Biolegend, Clone: 2H7, Cat#: 302304, RRID: AB_314252), CD2 (BD, Clone: RPA-2.10,Cat#: 555326), CD3 (BD, Clone: SK7, Cat#: 345763), CD16 (eBiosciences, Clone: eBioCB16, Cat#: 11-0168-41, RRID: AB_10804882), CD19 (eBiosciences, Clone: HIB19, Cat#: 11-0199-42, RRID: AB_10669461), CD235a (eBiosciences, Clone: HIR2, Cat#: 11-9987-82, RRID: AB_465477), CD66b (Biolegend, Clone: G10F5, Cat#: 305104, RRID: AB_314496), CD10 (Biolegend, Clone: HI10a, Cat#: 312208, RRID: AB_314919), CD127 (eBiosciences, Clone: RDR5, Cat#: 11-1278-42, RRID: AB_1907343), each FITC. JMML patient samples were also stained for CD96 expression (Laboratory of Martin Gramatzki, Clone: TH-111, Donkey anti-mouse PE (eBiosciences, Cat# 12-4010-82, RRID: AB_11063706) using the same panel described above with CD45RA BV650 (Biolegend, Clone: HI100, Cat# 304136, RRID: AB_2563653)

### Single cell clonogenic assays

Single HSPCs were sorted based on their immunophenotypic definitions into 96-well plates with 50 μL of methocult H4435 (Stemcell Technologies). Lineage output was assessed on day 7, day 10 and day 14 by evaluating the morphology of the colonies under direct light microscopy and individual colonies picked for cytospins, flow cytometry or resuspended in 10 μL of phosphate buffered saline (PBS), flash frozen and stored at −80 degrees for downstream mutational analysis. For targeting candidate JMML targets JQ1 (MedChem Express (Cat#: HY-13030), final concentration 250 nM), PD-0325901 (Cambridge Bioscience (Cat#: SM26), final concentrations 1uM or 5uM), Fasentin (Sigma-Aldrich, Cat#: F5557-5MG, final concentrations 5nM or 20nM) or equivalent DMSO vehicle control were added to the methocult prior to FACS sorting.

### Erythroid / megakaryocytic culture assay

Single cells were directly sorted into 96-well plates with erythroid/megakaryocyte bipotential medium for erythroid and megakaryocytic output (Psaila et al., 2016). Stemspan (Stemcell Technologies) was supplemented with TPO (100ng/ml), EPO (1 IU/ml), SCF (100ng/ml), IL-3 (10 ng/ml), IL-6 (10ng/ml) (all Peprotech), hu LDL (40 mg/ml) (Stemcell Technologies). Single cells were cultured at 37°C in 5% CO_2_ for 2 weeks and colony readout was performed by direct light microscopy and flow cytometry using CD41a APC (eBiosciences, Clone: HIP8, Cat#: 17-0419-42, RRID: AB_2573144), CD42b PE (eBiosciences, Clone: HIP1, Cat# 12-0429042, RRID: AB_10852864), CD11b FITC (eBiosciences,Clone: ICRF44, Cat#: 11-0118-42, RRID: AB_1582242), CD14 FITC (eBiosciences, Clone: 61D3, Cat#: 11-0149-41, RRID: AB_10597445), CD34 APC-eFlour780(eBiosciences, Clone: 4H11, Cat#: 47-0349-42, RRID: AB_2573956), and CD235a PerCp Cy5.5(Biolegend, Clone: HIR2, Cat#: 306614, RRID: AB_10683170).

### B-cell differentiation MS5 co-cultures

100 cells from relevant HSPC populations were sorted into cytokine-containing medium (AlphaMEM, Invitrogen), 10% heat-inactivated (batch tested) FBS, FLT3 (10 ng/ml), SCF (20ng/ml), IL2 (10 ng/ml), IL7 (5 ng/ml); all from (all Peprotech). Sorted cells were cultured on MS5 stroma cell lines (DSMZ, Cat#: ACC 441) for 4 weeks. Co-cultures were disaggregated by vigorous pipetting, passed through a 70 μm filter and replated on fresh MS5 cells every 4 days. Colony readout to assess the lymphoid output was performed by flow cytometry at week 2, 3 and 4 of the assay with CD11b FITC (eBiosciences,Clone: ICRF44, Cat#: 11-0118-42, RRID: AB_1582242), CD14 FITC (eBiosciences, Clone: 61D3, Cat#: 11-0149-41, RRID: AB_10597445), CD34 APC-eFlour 780 (eBiosciences, Clone: 4H11, Cat#: 47-0349-42, RRID: AB_2573956), CD235a PerCp Cy5.5 (Biolegend, Clone: HIR2, Cat#: 306614, RRID: AB_10683170), Human CD45 Alexa Flour 700 (Biolegend, Clone: HI130, Cat#: 304024, RRID: AB_493761), CD19 PE (BD, Clone: SJ25C1, Cat#: 345789), CD56 FITC (eBiosciences, Clone: TULY56, Cat#: 11-0566-42, RRID: AB_2572458), and CD33 FITC (BD, Clone: HIM3-4, Cat#: 555626). Lymphoid output for 2 control samples was assessed with Human CD45 Alexa Flour 700 (BD, Clone: 2D1, Cat#: 56-9459-42), CD19 APC (Biolegend, Clone: HIB19, Cat#: 302212), CD56 PE (eBiosciences, Clone: CMSSB, Cat#: 12-0567-41, RRID: AB_10598372), CD10 PE Cy7 (eBiosciences, Clone: ebioCB-CALLA, Cat#: 25-0106-42), CD34 PerCPCy5.5 (Biolegend, Clone: 581, Cat#: 343522), CD73 BV421 (Biolegend, Clone: AD2, Cat#: 344008), CD11b (eBiosciences,Clone: ICRF44, Cat#: 11-0118-42, RRID: AB_1582242), CD14 (eBiosciences, Clone: 61D3, Cat#: 11-0149-41, RRID: AB_10597445) and CD33 (BD, Clone: HIM3-4, Cat#: 555626) all FITC.

### Long-Term Culture-Initiating Cell (LTC-IC)

Stem cell potential was assessed in vitro using the Long-Term Culture-Initiating Cell Assay (LTC-IC) as previously described (Woll et al., 2014). Mouse stromal cell lines (M210B4 and SL/SL, sourced from Dr. Donna Hogge, Terry Fox Laboratory) were cultured for 7 days in RPMI supplemented with 10% FBS and DMEM supplemented with 15% FBS, respectively. Antibiotic selection was performed for an extra 7 days with hygromycin and G418, followed by irradiation 24 hours before FACS sorting. HSPCs from JMML patient samples and cord blood controls were isolated by FACS as described above and 100 cells/well were seeded in H5100 Myelocult medium (Stemcell Technologies) supplemented with 10^−6^ hydrocortisone 21-hemisuccinate (Stemcell Technologies) and co-cultured with SL/SL and M210B4 stroma cells. Sorted populations were cultured for 6 weeks, with half of the medium replaced once weekly. Individual wells were harvested after 6 weeks and transferred to Methocult H4435. Colony output was evaluated after 2 weeks under direct light microscopy.

### NSG xenotransplantation experiments

FACS purified HSPC populations were transplanted into NSG (NOD.Cg-Prkdc^scid^ II2rg^tm1Wjl^/SzJ) mice (Jackson Laboratory, Cat# 005557) of 10-14 weeks of age, following sublethal irradiation (split dose of 2.5 Gy delivered 4 hours apart). Cells were injected through tail vein injection 6-8 hours after the last radiation dose. Human cell reconstitution and lineage distribution was monitored by flow cytometry in the peripheral blood at 5, 10, 16, or 22 weeks post-transplant. For terminal readouts mice were sacrificed and flow cytometry analysis of BM, spleen and peripheral blood was performed in order to assess human cell reconstitution and immunophenotype of the engrafting population. BM MNC from primary transplants were cryopreserved and used for secondary transplantation. Peripheral blood was stained with Human CD45 Alexa Flour 700 (Biolegend, Clone: HI130, Cat#: 304024, RRID: AB_493761), Mouse CD45 PE-Texas Red (Invitrogen, Clone: 30-F11,Cat#: MCD4517, RRID: AB_10392557), CD19 PE (BD, Clone: SJ25C1, Cat#: 345789), CD34 APC-eFlour780 (eBiosciences, Clone: 4H11, Cat#: 47-0349-42, RRID: AB_2573956), CD235a PerCP Cy5.5 (Biolegend, Clone: HIR2, Cat#: 306614, RRID: AB_10683170), and CD15 (BD, Clone: MMA, Cat#: 347423), CD33 (BD, Clone: HIM3-4, Cat#: 555626), CD66b (Biolegend, Clone: G10F5, Cat#: 305104, RRID: AB_314496), CD11b (eBiosciences,Clone: ICRF44, Cat#: 11-0118-42, RRID: AB_1582242), CD14 (eBiosciences, Clone: 61D3, Cat#: 11-0149-41, RRID: AB_10597445) each FITC. Bone marrow was stained with CD34 APC-eFlour780 (eBiosciences, Clone: 4H11, Cat#: 47-0349-42, RRID: AB_2573956), CD38 PE-Texas Red (Invitrogen, Clone: HIT2, Cat#: MHCD3817, RRID: AB_10392545), CD90 BV421 (Biolegend, Clone: 5 E 10, Cat#: 328122, RRID: AB_2561420), CD45RA PE (eBiosciences, Clone: HI100, Cat#: 12-0458-41, RRID: AB_10717397), CD123 PE Cy7 (Biolegend, Clone: 6H6, Cat#: 306010, RRID: AB_493576), Mouse CD45 BV510 (Biolegend, Clone: 30-F11, Cat#: 103138, RRID: AB_2563061), Human CD45 AF700 (Biolegend, Clone: HI130, Cat#: 304024,RRID: AB_493761 and BD Clone: 2D1,Cat#: 56-9459-42), and CD8 (Biolegend, Clone: RPA-T8, Cat#: 301006, RRID: AB_314124), CD20 (Biolegend, Clone: 2H7, Cat#: 302304, RRID: AB_314252), CD2 (BD, Clone: RPA-2.10,Cat#: 555326), CD3 (BD, Clone: SK7, Cat#: 345763), CD16 (eBiosciences, Clone: eBioCB16, Cat#: 11-0168-41, RRID: AB_10804882), CD19 (eBiosciences, Clone: HIB19, Cat#: 11-0199-42, RRID: AB_10669461), CD235a (eBiosciences, Clone: HIR2, Cat#: 11-9987-82, RRID: AB_465477), CD66b (Biolegend, Clone: G10F5, Cat#: 305104, RRID: AB_314496), CD10 (Biolegend, Clone: HI10a, Cat#: 312208, RRID: AB_314919), CD127 (eBiosciences, Clone: RDR5, Cat#: 11-1278-42, RRID: AB_1907343) each FITC. Further details of the antibodies used can be found in the key resource table. NSG mice were bred at the Oxford Biomedical Services. All animal protocols were approved by the UK Home Office. For leukemia-free survival, mice were monitored for health and weight loss and euthanized if presenting with health concerns. Leukemia incidence was confirmed by presence of leukemic engraftment in peripheral blood >50%. Femurs preserved were fixed in neutral buffered formalin and embedded in paraffin for tissue sectioning and staining according to standard techniques.

### Bulk targeted next generation sequencing (NGS)

DNA was extracted from whole peripheral blood or BM using the Qiagen DNeasy blood and tissue kit (Qiagen, Manchester, UK). DNA was also extracted from hair and/or skin fibroblasts, to screen for germline mutations. A customized targeted NGS panel was designed, using the Illumina design studio (http://www.illumina.com/informatics/research/experimental-design/designstudio.html-design/designstudio.html), to screen for the most common mutations identified in this patient group. The NGS panel contained 32 genes with a total of 916 amplicons (covering 301 exons) (Table S 2). Coverage of each nucleotide within a target region was assessed for every sample on each sequencing run using in house customized software Covermi. Library preparation and sequencing was performed using the Illumina Truseq Custom Amplicon panel (TSCA) v1.5 kit (Illumina, Cat# WG-311-8007) and sequencing performed using the Illumina MiSeq with the v2 300 cycle reagent kit.

### Data analysis of the targeted NGS dataset

Data analysis was performed using in house bioinformatic pipelines at the Oxford Molecular Diagnostic center. In short, FASTQ files were processed by the MiSeq Reporter and sequences aligned to the reference genome (hg19), by the Smith-Waterman algorithm and Illumina Somatic Variant caller (1.1.0). Variants were annotated using Illumina VariantStudio (2.2) and were filtered using the following passing filters: quality >100 and consequence: missense, frameshift, stop gained, stop lost, initiator codon, in frame insertion, in-frame deletion, splice filters. Variants were first verified by visual check by using the Interactive Genomics Viewer (IGV, 2.3.40) and confirmed by either Sanger sequencing or PCR-based mutational analysis performed in the Great Ormond Street Hospital diagnostic lab.

### Mutational analysis of colonies and single cells

Mutations were traced back to the cell of origin by mutational analysis of colonies that arise from single cells sorted from each population. Patient-specific primers were designed for downstream amplification of the region of interest using the Primer blast tool website (https://www.ncbi.nlm.nih.gov/tools/primer-blast/). The sequences of primers used can be found in Table S3. For every individual cell sorted for genotyping, all index sort data of the MFI for CD34, CD38, CD90, CD45RA and CD123 were recorded. This allowed the immunophenotypic identification of every cell processed and mapping of the JMML mutations onto the hematopoietic hierarchy.

Colonies derived from sorted single cells from each population were picked and resuspended in 10 µL PBS and frozen at −80^0^C for downstream analysis. Samples were thawed and lysed using 15 µL of a custom lysis buffer containing 10% Proteinase K (Qiagen) and 0.4% Triton X. Samples were incubated at 56°C for 10 minutes and 75°C for 20 minutes to release genomic DNA and inactivate the Proteinase K, respectively. Regions of interest were amplified using a 3-step nested PCR method with FastStart High Fidelity enzyme (Roche). For the first PCR, 3.6 µL of the lysate was amplified with patient specific primers covering the regions of interest with a targeted length of ∼300bp amplicons. For the second PCR step, 1 µL of the PCR1 product was amplified with nested target specific primers generating ∼ 200bp amplicons. Primers for PCR 2 were attached to Access Array adaptors (Fluidigm; Forward adaptor: ACACTGACGACATGGTTCTACA; Reverse adaptor: TACGGTAGCAGAGACTTGGTCT). Lastly, for the third PCR step, Illumina compatible adaptors containing single-direction indexes (Access ArrayTM Barcode Library for Illumina Sequencers-384, Single Direction, Fluidigm) were attached to pre-amplified amplicons from PCR2. This final PCR product was purified with Ampure XP beads (0.8: to 1 ratio beads to product), quantified using Quant-iT Picogreen (Thermo Fisher Scientific), pooled and sequenced on the Illumina MiSeq with 150 bp paired-end reads.

### Single cell colony genotyping analysis

Reads were aligned to the human genome (GRCh37/hg19) using STAR with default settings (Cold Spring Harbor Laboratory, version 2.4.2a, RRID:SCR_004463). Variant calling was performed using mpileup (samtools, Wellcome Trust Sanger Institute, version 1.1, RRID:SCR_002105, options -- minBQ 30, --count-orphans, --ignore overlaps) and results were summarized with a custom Python script. Downstream analysis of the variants and annotation of the mutant cells was performed with a custom R script. Each individual amplicon was analyzed and those with coverage < 30 reads were excluded from the downstream analysis. Samples in which one of the amplicons was not detected and samples with allelic dropout (assessed using SNPs) in three or more amplicons were also excluded from further analysis. For each sequencing run, normal controls were also screened for the variants of interest in order to ensure no false positive samples were present. None of the mutations were detected in any of the control samples in any of the sequencing runs. Amplicons covering common heterozygous SNPs for each patient were used to calculate the allelic dropout rate of the assay, as described at Supplementary Figure 3.

### Bulk RNA sequencing analysis

Data analysis was performed with in house bioinformatics pipelines. Briefly, RNA-sequencing reads from bulk population were aligned to the human genome (GRCh38) using STAR (Cold Spring Harbor Laboratory,version 2.4.2a, RRID:SCR_004463). Aligned reads were counted using featureCounts (Walter+Eliza Hall Institute of Medical Research, version 1.4.5-p1, RRID:SCR_012919) summarized to gene identifiers from the GRCh38 Ensembl transcriptome. Differential gene expression was done using the DESeq2 package (Bioconductor, 1.16.1, RRID:SCR_015687) in R (CRAN, 3.4.1). Plotting expression values of genes was done by calculating transcripts per million (TPM) and plotted in R. Gene-set enrichment analysis was performed using GSEA software (Broad Institute,v 2.20, RRID:SCR_003199) against Hallmark and curated HSC and cell cycle gene sets obtained through MSigDB (Broad Institute, RRID:SCR_003199) using the default settings. Heatmaps were generated using the package pheatmap (1.0.8).

### Single cell RNA sequencing

Reads from 10x genomics single cell RNA-sequencing were aligned and counted using cell ranger (2.0.0) using the hg19 human genome. Unique molecular identifier (UMI) counts where analyzed using Seurat (Satija Lab, v 2.0.1). Genes were filtered by minimum number of cells expressing = 10, greater than 0.15 mean UMI counts, less than 4 mean UMI counts, and greater than 0.5 dispersion. Cells were filtered by greater than 1000 genes detected, less that 4000 genes detected, and less than 5% mitochondrial reads. After filtering 1,127 variable genes and 17,547 cells were used for analysis. tSNE analysis was performed after selecting the most variable principle components by use of the jackStrawPlot feature in Seurat. Clusters were identified by the k-shared nearest neighbor (SSN) approach FindClusters. Figures for single cell analysis were generated in R. Top differentially expressed genes from SSN identified clusters were calculated using FindAllMarkers function in Seurat.

### QUANTIFICATION AND STATISTICAL ANALYSIS

T-tests. Mann Whitney tests and ANOVA followed by multiple comparisons testing were used to compare experimental groups as indicated in the figure legends. Statistical analyses were performed using GraphPad Prism v7.00 or R v3.4.1. Survival curves were compared using the log-rank test.

### SOFTWARE AVAILABILITY

R and Python scripts used for the analysis are available upon request.

## References

1. F. Locatelli, C. M. Niemeyer, How I treat juvenile myelomonocytic leukemia. Blood 125, 1083–1090 (2015); published online EpubFeb 12 (10.1182/blood-2014-08-550483).

2. A. Caye, M. Strullu, F. Guidez, B. Cassinat, S. Gazal, O. Fenneteau, E. Lainey, K. Nouri, S. Nakhaei-Rad, R. Dvorsky, J. Lachenaud, S. Pereira, J. Vivent, E. Verger, D. Vidaud, C. Galambrun, C. Picard, A. Petit, A. Contet, M. Poiree, N. Sirvent, F. Mechinaud, D. Adjaoud, C. Paillard, B. Nelken, Y. Reguerre, Y. Bertrand, D. Haussinger, J. H. Dalle, M. R. Ahmadian, A. Baruchel, C. Chomienne, H. Cave, Juvenile myelomonocytic leukemia displays mutations in components of the RAS pathway and the PRC2 network. Nat Genet 47, 1334–1340 (2015); published online EpubNov (10.1038/ng.3420).

3. E. Stieglitz, A. N. Taylor-Weiner, T. Y. Chang, L. C. Gelston, Y. D. Wang, T. Mazor, E. Esquivel, A. Yu, S. Seepo, S. Olsen, M. Rosenberg, S. L. Archambeault, G. Abusin, K. Beckman, P. A. Brown, M. Briones, B. Carcamo, T. Cooper, G. V. Dahl, P. D. Emanuel, M. N. Fluchel, R. K. Goyal, R. J. Hayashi, J. Hitzler, C. Hugge, Y. L. Liu, Y. H. Messinger, D. H. Mahoney, Jr., P. Monteleone, E. R. Nemecek, P. A. Roehrs, R. J. Schore, K. C. Stine, C. M. Takemoto, J. A. Toretsky, J. F. Costello, A. B. Olshen, C. Stewart, Y. Li, J. Ma, R. B. Gerbing, T. A. Alonzo, G. Getz, T. Gruber, T. Golub, K. Stegmaier, M. L. Loh, The genomic landscape of juvenile myelomonocytic leukemia. Nat Genet 47, 1326–1333 (2015); published online EpubNov (10.1038/ng.3400).

4. J. A. Magee, E. Piskounova, S. J. Morrison, Cancer stem cells: impact, heterogeneity, and uncertainty. Cancer Cell 21, 283–296 (2012); published online EpubMar 20 (10.1016/j.ccr.2012.03.003).

5. A. J. Mead, A. Mullally, Myeloproliferative neoplasm stem cells. Blood 129, 1607–1616 (2017); published online EpubMar 23 (10.1182/blood-2016-10-696005).

6. P. Gallipoli, S. A. Abraham, T. L. Holyoake, Hurdles toward a cure for CML: the CML stem cell. Hematol Oncol Clin North Am 25, 951–966, v (2011); published online EpubOct (10.1016/j.hoc.2011.09.001).

7. P. S. Woll, U. Kjallquist, O. Chowdhury, H. Doolittle, D. C. Wedge, S. Thongjuea, R. Erlandsson, M. Ngara, K. Anderson, Q. Deng, A. J. Mead, L. Stenson, A. Giustacchini, S. Duarte, E. Giannoulatou, S. Taylor, M. Karimi, C. Scharenberg, T. Mortera-Blanco, I. C. Macaulay, S. A. Clark, I. Dybedal, D. Josefsen, P. Fenaux, P. Hokland, M. S. Holm, M. Cazzola, L. Malcovati, S. Tauro, D. Bowen, J. Boultwood, A. Pellagatti, J. E. Pimanda, A. Unnikrishnan, P. Vyas, G. Gohring, B. Schlegelberger, M. Tobiasson, G. Kvalheim, S. N. Constantinescu, C. Nerlov, L. Nilsson, P. J. Campbell, R. Sandberg, E. Papaemmanuil, E. Hellstrom-Lindberg, S. Linnarsson, S. E. Jacobsen, Myelodysplastic syndromes are propagated by rare and distinct human cancer stem cells in vivo. Cancer Cell 25, 794–808 (2014); published online EpubJun 16 (10.1016/j.ccr.2014.03.036).

8. D. Thomas, R. Majeti, Biology and relevance of human acute myeloid leukemia stem cells. Blood 129, 1577–1585 (2017); published online EpubMar 23 (10.1182/blood-2016-10-696054).

9. C. le Viseur, M. Hotfilder, S. Bomken, K. Wilson, S. Rottgers, A. Schrauder, A. Rosemann, J. Irving, R. W. Stam, L. D. Shultz, J. Harbott, H. Jurgens, M. Schrappe, R. Pieters, J. Vormoor, In childhood acute lymphoblastic leukemia, blasts at different stages of immunophenotypic maturation have stem cell properties. Cancer Cell 14, 47–58 (2008); published online EpubJul 8 (10.1016/j.ccr.2008.05.015).

10. M. Jan, T. M. Snyder, M. R. Corces-Zimmerman, P. Vyas, I. L. Weissman, S. R. Quake, R. Majeti, Clonal evolution of preleukemic hematopoietic stem cells precedes human acute myeloid leukemia. Sci Transl Med 4, 149ra118 (2012); published online EpubAug 29 (10.1126/scitranslmed.3004315).

11. N. J. Palpant, Y. Wang, B. Hadland, R. J. Zaunbrecher, M. Redd, D. Jones, L. Pabon, R. Jain, J. Epstein, W. L. Ruzzo, Y. Zheng, I. Bernstein, A. Margolin, C. E. Murry, Chromatin and Transcriptional Analysis of Mesoderm Progenitor Cells Identifies HOPX as a Regulator of Primitive Hematopoiesis. Cell Rep 20, 1597–1608 (2017); published online EpubAug 15 (10.1016/j.celrep.2017.07.067).

12. L. L. Remsing Rix, U. Rix, J. Colinge, O. Hantschel, K. L. Bennett, T. Stranzl, A. Muller, C. Baumgartner, P. Valent, M. Augustin, J. H. Till, G. Superti-Furga, Global target profile of the kinase inhibitor bosutinib in primary chronic myeloid leukemia cells. Leukemia 23, 477–485 (2009); published online EpubMar (10.1038/leu.2008.334).

13. C. A. G. Booth, N. Barkas, W. H. Neo, H. Boukarabila, E. J. Soilleux, G. Giotopoulos, N. Farnoud, A. Giustacchini, N. Ashley, J. Carrelha, L. Jamieson, D. Atkinson, T. Bouriez-Jones, R. K. Prinjha, T. A. Milne, D. T. Teachey, E. Papaemmanuil, B. J. P. Huntly, S. E. W. Jacobsen, A. J. Mead, Ezh2 and Runx1 Mutations Collaborate to Initiate Lympho-Myeloid Leukemia in Early Thymic Progenitors. Cancer Cell 33, 274–291 e278 (2018); published online EpubFeb 12 (10.1016/j.ccell.2018.01.006).

14. T. De Raedt, E. Beert, E. Pasmant, A. Luscan, H. Brems, N. Ortonne, K. Helin, J. L. Hornick, V. Mautner, H. Kehrer-Sawatzki, W. Clapp, J. Bradner, M. Vidaud, M. Upadhyaya, E. Legius, K. Cichowski, PRC2 loss amplifies Ras-driven transcription and confers sensitivity to BRD4-based therapies. Nature 514, 247–251 (2014); published online EpubOct 9 (10.1038/nature13561).

15. N. Hosen, C. Y. Park, N. Tatsumi, Y. Oji, H. Sugiyama, M. Gramatzki, A. M. Krensky, I. Weissman, CD96 is a leukemic stem cell-specific marker in human acute myeloid leukemia. Proc Natl Acad Sci U S A 104, 11008–11013 (2007); published online EpubJun 26 (10.1073/pnas.0704271104).

16. M. Dimitriou, P. S. Woll, T. Mortera-Blanco, M. Karimi, D. C. Wedge, H. Doolittle, I. Douagi, E. Papaemmanuil, S. E. W. Jacobsen, E. Hellstrom-Lindberg, Perturbed hematopoietic stem and progenitor cell hierarchy in myelodysplastic syndromes patients with monosomy 7 as the sole cytogenetic abnormality. Oncotarget 7, 72685–72698 (2016); published online EpubNov 8 (10.18632/oncotarget.12234).

17. S. L. Prashad, V. Calvanese, C. Y. Yao, J. Kaiser, Y. Wang, R. Sasidharan, G. Crooks, M. Magnusson, H. K. Mikkola, GPI-80 defines self-renewal ability in hematopoietic stem cells during human development. Cell Stem Cell 16, 80–87 (2015); published online EpubJan 8 (10.1016/j.stem.2014.10.020).

18. B. J. Povinelli, A. Rodriguez-Meira, A. J. Mead, Single cell analysis of normal and leukemic hematopoiesis. Mol Aspects Med 59, 85–94 (2018); published online EpubFeb (10.1016/j.mam.2017.08.006).

19. A. Giustacchini, S. Thongjuea, N. Barkas, P. S. Woll, B. J. Povinelli, C. A. G. Booth, P. Sopp, R. Norfo, A. Rodriguez-Meira, N. Ashley, L. Jamieson, P. Vyas, K. Anderson, A. Segerstolpe, H. Qian, U. Olsson-Stromberg, S. Mustjoki, R. Sandberg, S. E. W. Jacobsen, A. J. Mead, Single-cell transcriptomics uncovers distinct molecular signatures of stem cells in chronic myeloid leukemia. Nat Med 23, 692–702 (2017); published online EpubJun (10.1038/nm.4336).

20. S. Picelli, O. R. Faridani, A. K. Bjorklund, G. Winberg, S. Sagasser, R. Sandberg, Full-length RNA-seq from single cells using Smart-seq2. Nature protocols 9, 171–181 (2014); published online EpubJan (10.1038/nprot.2014.006).

